# Aberrant astrocyte protein secretion contributes to altered neuronal development in diverse disorders

**DOI:** 10.1101/2020.02.17.939991

**Authors:** Alison L.M. Caldwell, Jolene K. Diedrich, Maxim N. Shokhirev, Nicola J. Allen

**Affiliations:** Molecular Neurobiology Laboratory, Salk Institute for Biological Studies, 10010 North Torrey Pines Rd, La Jolla, CA, 92037, USA; Mass Spectrometry Core, Salk Institute for Biological Studies, 10010 North Torrey Pines Rd, La Jolla, CA, 92037, USA; Razavi Newman Integrative Genomics and Bioinformatics Core, Salk Institute for Biological Studies, 10010 North Torrey Pines Rd, La Jolla, CA, 92037, USA; Neurosciences Graduate Program University of California San Diego La Jolla, CA, 92093 USA

## Abstract

Astrocytes negatively impact neuronal development in many neurodevelopmental disorders (NDs), however how they do this, and if mechanisms are shared across disorders, is not known. We developed an in vitro system to ask how astrocyte protein secretion and gene expression change in three genetic NDs. We identified disorder specific changes, and changes common to all disorders. ND astrocytes increase release of Igfbp2, a secreted inhibitor of IGF. IGF rescues neuronal deficits in many NDs, and we found blocking Igfbp2 partially rescues inhibitory effects of Rett Syndrome astrocytes, suggesting increased astrocyte Igfbp2 contributes to decreased IGF signaling in NDs. We identified increased BMP signaling in ND astrocytes is upstream of protein secretion changes, including Igfbp2, and blocking BMP signaling in Fragile X Syndrome astrocytes reverses inhibitory effects on neurite outgrowth. We provide a resource of astrocyte secreted proteins in health and NDs, and identify novel targets for intervention in diverse NDs.

## Introduction

The formation of neuronal circuits is a complex process requiring coordinated signaling between multiple cell types in the developing brain. In the rodent cortex this begins embryonically with neuronal generation, and proceeds throughout the first month of postnatal life with neuronal migration, axon and dendrite growth, synapse formation and maturation, and elimination of inappropriate synapses^1^. Perturbation at any of these stages, as occurs in neurodevelopmental disorders (ND), can have life-long consequences for brain function. Despite being caused by diverse genetic mutations, many different NDs have over-lapping phenotypic neuronal alterations, including stunting of axonal and dendritic arbor growth, deficits in neuronal synapse formation, and immature dendritic spines^2^. For example in Rett Syndrome (RTT; Mecp2 loss of function) there is a reduction in overall dendritic arbor complexity and a decrease in spine density^3, 4^; in Fragile X Syndrome (FXS; Fmr1 loss of function) dendritic outgrowth is stunted, and there is an increase in thin, immature spines^5, 6^; in Down Syndrome (DS; trisomy chromosome 21), there is a reduction in spine and synapse number, and an increase in immature spines^7^. The phenotypic overlap in these NDs suggests they may share commonly altered molecular mechanisms, raising the possibility of identifying new targets for therapy development across diverse NDs.

The majority of ND research has focused on intrinsic changes within neurons, and has identified some therapeutic targets. However, emerging research has identified that not all defects in NDs are intrinsic to neurons and that alterations to glial cell function are playing a role^8^. Astrocytes are an abundant class of glial cell, and in the healthy brain support all stages of neuronal development predominantly through secreted factors^9^. For example, wild-type (WT) neurons grown in vitro in the absence of astrocytes show poor survival, stunted neurite outgrowth and defects in synapse formation, whereas the addition of WT astrocytes or astrocyte secreted proteins (ACM) is sufficient to reverse these deficits and promote neuronal development^10–12^. Conversely, WT neurons cultured with astrocytes or ACM from FXS, RTT or DS fail to develop normally, exhibiting stunted neurite outgrowth, decreased synapse formation and immature dendritic spines^8, 13–16^. This demonstrates that astrocytes are contributing to aberrant neuronal development observed in these disorders through an alteration in the release of secreted factors.

To determine how ND astrocytes inhibit neuronal development, we developed an in vitro system to identify protein secretion and gene expression of cortical astrocytes from WT and ND mouse models (FXS, RTT, DS). We chose these three disorders due to the overlapping phenotype induced in WT neurons by astrocytes from each disorder. We developed an immunopanning (IP) procedure to prospectively isolate age and region-matched astrocytes and neurons from the postnatal day (P) 7 mouse cortex, and maintained them in vitro in defined serum-free media. The use of age and region matched neurons and astrocytes is particularly important given that astrocytes from different brain regions have significantly different gene expression, as well as locally specialized properties^17, 18^. Using IP to isolate astrocytes has several advantages over the classical astrocyte culture method^19^. IP allows astrocyte isolation at older ages e.g. P7 used here, when they are actively regulating neurite and synapse development; in the classical procedure astrocytes are isolated at P0-1 when the cells are immature and represent progenitors. Culturing IP astrocytes in defined serum-free media maintains their gene expression profile so it closely matches that of acutely isolated astrocytes; in the classical procedure astrocytes are cultured in serum which profoundly alters their properties^20^. For example, serum induces genes associated with inflammatory reactive astrocytes, confounding results by making all astrocytes reactive even in the absence of disease.

Using the IP system we have generated a resource of the protein secretion and gene expression profiles of WT and ND astrocytes from the developing cortex. We used this resource to identify how astrocytes from FXS, RTT and DS differ in protein secretion from WT, finding unique changes for each disorder, as well as 88 proteins that are significantly increased in secretion across all 3 NDs. Interestingly, although gene expression is altered between WT and ND astrocytes, there is little overlap in protein secretion and gene expression changes, highlighting the importance of a proteomics approach when studying secreted proteins^21^. We validated that two of the proteins upregulated in secretion from ND astrocytes, Igfbp2 and BMP6, are upregulated in vivo in mouse models of ND, and have negative effects on neuronal development that can be rescued by antagonizing their function. Altered IGF signaling is implicated in multiple NDs, and treatment of RTT and FXS mice with IGF1 improves behavioral and physiological symptoms^22–24^. Igfbp2 is a secreted inhibitor of IGF signaling, and given the positive effects of IGF1 in NDs we hypothesize that increased Igfbp2 release from astrocytes in NDs is inhibiting endogenous IGF signaling. BMP6 is a member of the transforming growth factor beta superfamily, and BMPs regulate astrocyte maturation^25^. We found that BMP signaling in astrocytes is upstream of many of the protein secretion changes in ND astrocytes, including Igfbp2. This work has identified how altered astrocyte protein release contributes to aberrant neuronal development in diverse NDs. This provides new pathways for therapeutic targeting in these disorders, as well as in sporadic NDs that have shared phenotypic neuronal alterations.

## Results

### Immunopanned astrocyte and neuron cultures for the study of NDs

To enable the study of astrocytes from mouse models of NDs, we developed an immunopanning (IP) procedure to isolate astrocytes from the cortex. We modified the protocol previously developed for rat astrocytes by changing the antibody used for positive selection to one that recognizes mouse astrocytes (for detailed procedure see Methods)^20, 26^. The cerebral cortices from P7 mice were digested with papain to produce a single cell suspension, which was passed over antibody-coated plates to deplete irrelevant cell types (microglia, endothelial cells, oligodendrocyte precursor cells (OPCs)). Positive selection of astrocytes was performed using an antibody against astrocyte cell surface protein 2 (ACSA2; recognizes Atp1b2) (Figure 1a)^27, 28^. Isolated astrocytes were maintained in serum-free media with the addition of the growth factor HB-EGF (heparin binding epidermal growth factor) to support cell survival. The absence of serum is essential to prevent the induction of genes associated with reactive astrocytes^20^. qRT-PCR analysis of astrocyte mRNA determined that IP astrocytes are highly enriched for astrocyte markers and depleted for other cell-type markers (neuron, microglia, OPC, fibroblast) (Figure 1b). Further, immunostaining cultures for cell-type specific markers showed the majority of cells are positive for the astrocyte marker GLAST (Slc1a3) (Figure 1c), with neurons and microglia not detected, and OPCs rarely detected (Figure S1a,b). Astrocytes were cultured for 5 to 7 days until confluent, then switched to a protein-free conditioning media for a further 5 days in order to collect astrocyte-secreted proteins, referred to as astrocyte conditioned media (ACM). The use of protein free media is essential to enable the detection of low abundance proteins secreted by astrocytes using mass spectrometry analysis.

**Figure 1.**
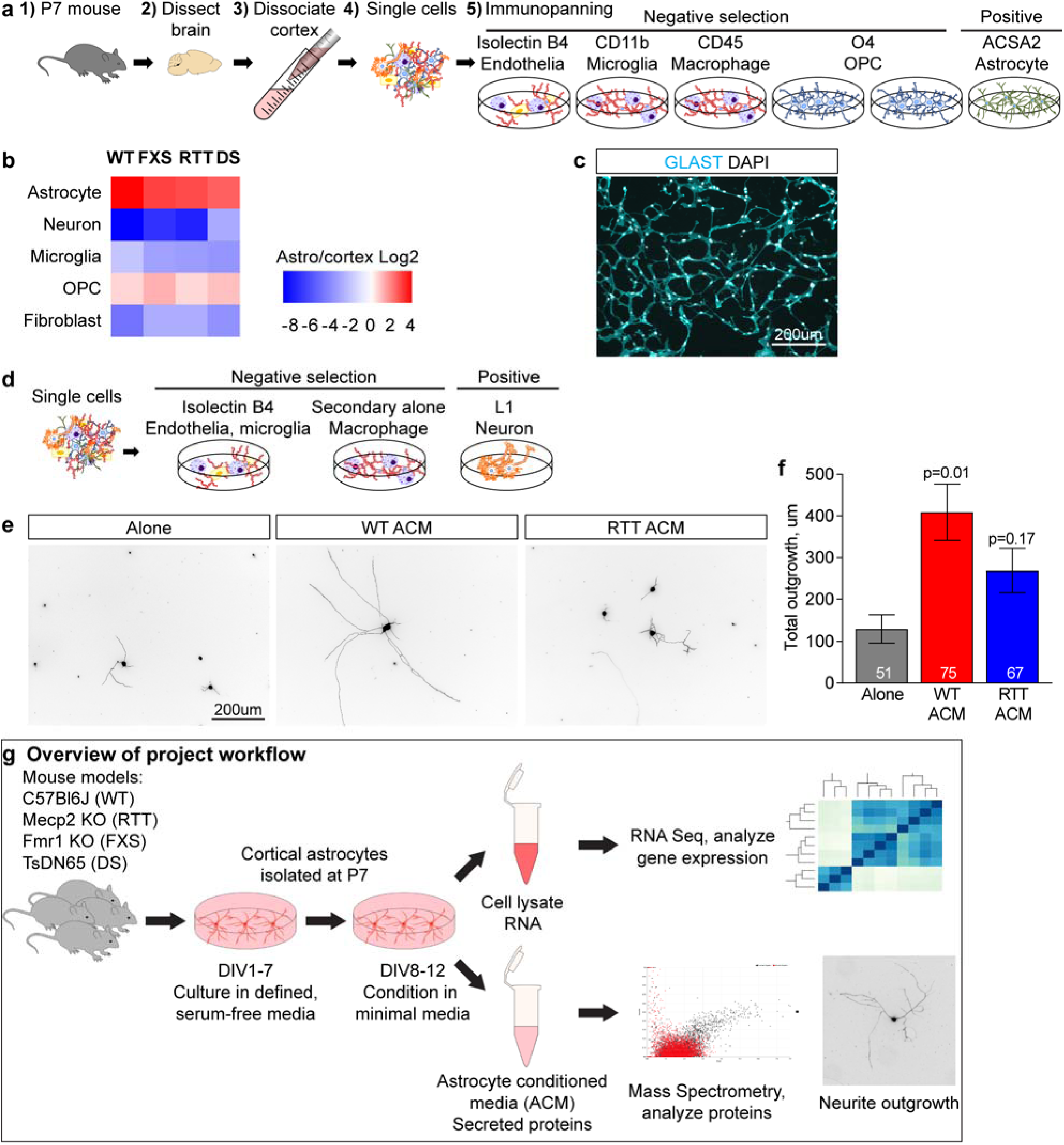
Immunopanned astrocyte and neuron cultures for the study of NDs. **a.** Schematic of the procedure: P7 mouse cortex is digested with papain to produce a single cell suspension, which undergoes a series of negative selection steps to deplete unwanted cells (endothelia, microglia, oligodendrocytes), followed by positive selection for astrocytes using an antibody against ACSA2. **b.** qRT-PCR for cell-type markers from mRNA collected from IP astrocytes compared to P7 mouse cortex demonstrates enrichment for astrocytes (Gfap), a depletion of neurons (Syt1), microglia (Cd68), fibroblasts (Fgfr4), and a decrease in oligodendrocyte precursor cells (OPCs; Cspg4) in WT and ND IP astrocyte cultures. N=6 cultures per genotype. **c.** Immunostaining IP-astrocyte cultures for astrocyte marker GLAST (cyan, Slc1a3) and nuclei (white, DAPI) demonstrates the majority of cells express this protein. **d-f.** WT ACM supports WT neurite outgrowth, whereas RTT ACM does not. **d.** Schematic of the immunopanning procedure to isolate cortical neurons: P7 mouse cortex is digested with papain to produce a single cell suspension, which undergoes a series of negative selection steps to deplete unwanted cells (endothelia, microglia), followed by positive selection for neurons using an antibody against NCAM-L1. **e-f.** Culturing WT cortical neurons for 48 hours with WT astrocyte conditioned media (ACM) increases neurite outgrowth, whereas RTT ACM does not. **e.** Example images, neurons immunostained with MAP2 and tau, merged image shown. **f.** Quantification of total neurite outgrowth (dendrite + axon). Example experiment shown, experiment repeated 3 times with same result. Bar graph represents mean ± s.e.m. Number in bar = neurons analyzed. Statistics one-way ANOVA on ranks, p value compared to neurons alone. **g.** Overview of project workflow. See also Figure S1.

First we asked if ACM from IP astrocytes has the same effects on neurite outgrowth as those reported in the literature, i.e. WT ACM supports neurite outgrowth, and ND ACM inhibits it^13^. To do this we modified the immunopanning protocol to isolate age and region matched neurons from the WT P7 mouse cortex, using an antibody against the neuronal protein NCAM-L1 (Figure 1d)^29^. Cortical neurons were plated in serum free medium (alone condition) or with ACM (3μg/ml) added at the time of plating, cultured for 48 hours, and neurite outgrowth assessed by immunostaining neurons for Map2 (dendrites) and Tau (axons) (Figure 1e,f; S1c-e). We chose 48 hours as this time period has been used in multiple previous studies investigating the effects of ND astrocytes on neurite development^22, 30^. We found WT cortical neurons grown in isolation show minimal neurite outgrowth (total outgrowth = Map2+Tau), and addition of WT ACM significantly increases total neurite outgrowth. Treating WT neurons with RTT ACM does not induce a significant increase in neurite outgrowth compared to untreated neurons (Alone 130 ± 34µm, WT ACM 422 ± 68µm, RTT ACM 269 ± 53µm; Figure 1e,f; S1c-e). We analyzed a number of additional parameters in addition to total neurite outgrowth, including longest neurite, mean process length and number of branches, and found the same effect for all measures, i.e. WT ACM supports neurite development and RTT ACM does not (Fig S1e). Based on this in all further experiments we used total neurite outgrowth (Map2+Tau) to compare the effects of WT and ND ACM on WT neuron development.

The cortical neuron outgrowth experiments demonstrate that ACM from RTT astrocytes does not support neurite outgrowth, whereas WT ACM does, indicating that a difference in the composition of proteins secreted from the astrocytes is responsible. To analyze what the differences are, and to see if they are shared with astrocytes from other NDs, we compared protein secretion and gene expression from astrocytes from 3 mouse models of NDs to strain matched WT control: 1) RTT, Mecp2 KO; 2) FXS, Fmr1 KO; 3) DS, Ts65Dn Down syndrome transgenic with one copy of the duplicated chromosome; 4) WT control, C57Bl6J (Figure 1g)^31–33^. Although RTT and FXS are predominantly seen in females with one altered copy of the gene we chose to analyze astrocyte cultures from KO mice. This is because heterozygous cultures would be a mixture of WT and KO cells due to Mecp2 and Fmr1 being present on the X chromosome, and random inactivation of the X chromosome occurring, which would mask effects of altered protein secretion into the media. To analyze protein secretion we used mass spectrometry to identify proteins present in ACM from 6 separate cultures per genotype. To focus on proteins that are robustly produced we filtered data for proteins that are present at 0.01% or above of total protein in one of the genotypes. This identified 1235 unique proteins in WT ACM (Table S1). Patternlab software was used to determine if proteins are present at significantly different levels between ACM from WT and each ND, with significance set at p<0.05, abundance ≥0.01%, fold change between WT and ND ≥1.5. For RNA sequencing analysis of gene expression, 6 separate cultures of WT, FXS and RTT, along with 4 separate cultures of DS were analyzed. For WT astrocytes, 12315 genes are present at FPKM>1 (Table S4). Significance for differential expression between ND and WT was calculated in DESeq2 and defined as p<0.05 adjusted for multiple comparisons, along with expression ≥FPKM 1 and fold change between ND and WT ≥1.5.

### Immunopanned astrocytes reproduce known in vivo alterations to ND astrocyte function

We first asked if IP astrocytes resemble in vivo astrocytes by analyzing expression levels of genes associated with astrocyte function. This includes astrocyte markers GFAP, Aqp4 and Aldh1l1, all of which are expressed at high levels by IP astrocytes (Figure 2a,b)^34^. We found no significant differences in marker gene expression between WT and ND, except the calcium binding protein S100b which is significantly more highly expressed by DS astrocytes (Log2 fold change vs WT (LFC) 1.10, p<0.0001), as previously reported in DS patient blood samples^35^. This shows no large changes in cellular identity occur in ND astrocytes. We next examined ion channels and transporters associated with astrocyte function, including the potassium channel Kir4.1 (Kcnj10), the glutamate uptake transporters Glt1 (Slc1a2) and Glast (Slc1a3), and metabolic enzymes including Ldha (lactate dehydrogenase) and Srebf1 (lipid synthesis). These are expressed at high levels by IP astrocytes and at similar levels across WT and ND, except Glt1 which is decreased in FXS (LFC −0.79, p=0.002), and the GABA uptake transporter Slc6a11 which is decreased in both FXS (LFC −0.65, p=0.02) and DS (LFC −0.98, p=0.003). Decreased astrocyte Glt1 has been reported in vivo in FXS mice, however astrocytes cultured from FXS mice using the traditional method do not show this difference^36, 37^. As the Glt1 decrease is still present in IP FXS astrocytes, this validates this system for maintaining pathological astrocyte properties in vitro, and provides a means for their study. We further asked if expression of neurotransmitter receptors is altered between astrocytes from ND and WT, and found that metabotropic glutamate receptors Grm3 and Grm5 are downregulated in FXS astrocytes (Grm3: LFC −1.04, p=0.007; Grm5: LFC −1.98, p=0.002), whilst Grm5 is decreased in DS (LFC −2.13, p<0.0001). As with Glt1, the decrease in Grm5 has previously been reported in FXS astrocytes in vivo^37^.

**Figure 2.**
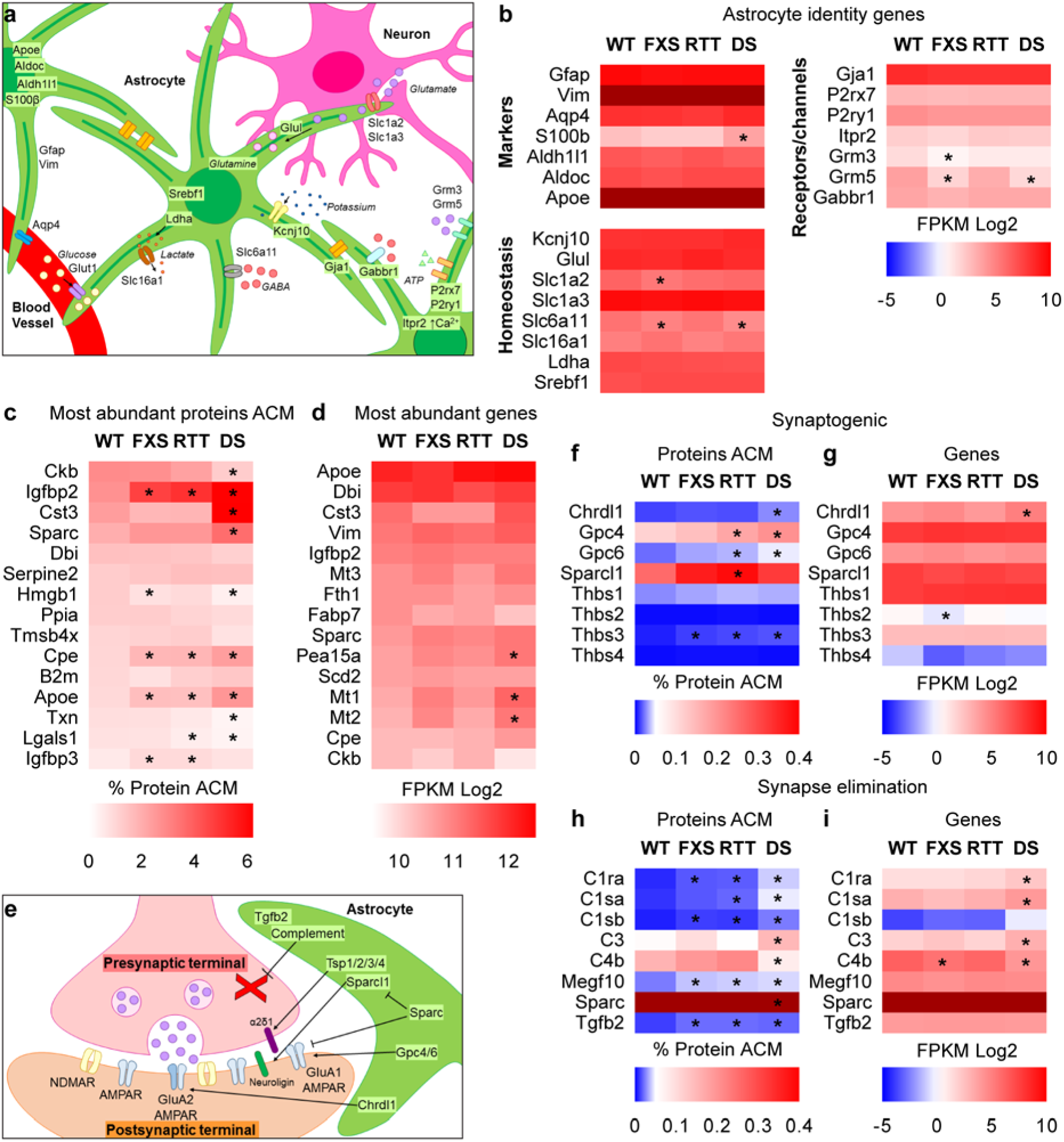
Immunopanned astrocytes reproduce known alterations to ND astrocyte function. **a,b.** WT and ND IP astrocytes express many known astrocyte markers at high levels. **a.** Astrocytes (green) express cell-specific markers that determine their cellular identity (top left astrocyte), contact blood vessels and neuronal synapses to engage in metabolism and homeostatic functions (center astrocyte), and bind and respond to neurotransmitters released by neurons (lower right astrocyte). **b.** Heatmap shows few differences between ND and WT expression of astrocyte identity and function markers. **c.** Heatmap of most abundant proteins, ranked by level in WT ACM. **d.** Heatmap of most abundant mRNA, ranked by level in WT astrocytes. **e-i.** Schematic of the tripartite synapse (**e**) displaying astrocyte-secreted proteins important for regulating synapse formation and function. **f-i.** Heatmaps of secreted synaptogenic proteins in ACM (**f**) and expression of synaptogenic genes (**g**), as well as abundance of synapse eliminating proteins in ACM (**h**) and corresponding expression of synapse elimination genes (**i**). Proteomics, N=6 cultures per genotype, *p<0.05, abundance >0.01%, fold change between WT and ND ≥1.5. RNASeq, N=6 cultures WT, RTT, FXS; 4 cultures DS, * adjusted p<0.05, FPKM>1, fold change between ND and WT ≥1.5. See also Figure S2; Table S1, S4.

A hypothesis for why ND astrocytes have a negative impact on neuronal development is that they are in a reactive state and produce damaging inflammatory mediators^38, 39^. We compared the fold-change in gene expression between each ND and WT for a panel of reactive astrocyte genes, and found no consistent alterations (Figure S2a)^40^. While this does not rule out that ND astrocytes show reactive properties in vivo, this means IP astrocytes allow study of non-reactive changes to astrocytes that occur in ND, and identification of novel non-reactive targets.

We next determined the most abundant proteins secreted by, and genes expressed by, WT IP astrocytes (Figure 2c,d). Both lists include well known astrocyte-secreted proteins such as Dbi (diazepam binding inhibitor, a modulator of GABA receptors), Apoe (apolipoprotein E, a component of astrocyte lipid particles) and Sparc (negative regulator of synapse formation)^34, 41^. In the top 15 genes there are only 3 significant differences between WT and ND, all in DS astrocytes: an upregulation in Mt1 and Mt2 (metallothionein family members, zinc binding proteins) (as previously reported for Mt3 in DS^42^) (Mt1: LFC 0.79, p=0.04; Mt2: LFC 0.81, p=0.03) and an upregulation in Pea15a (Proliferation and apoptosis adaptor protein 15) (LFC 0.58, p=0.0003). There are more significant differences in protein secretion, of note an increase in Igfbp2 (insulin like growth factor binding protein 2, a secreted binding protein for insulin growth factor (IGF); Fold change vs WT (FC) FXS: 1.72, p=0.004; RTT: FC 1.87, p<0.0001; DS: FC 2.31, p<0.0001), Cpe (carboxypeptidase E, a component of the regulated secretory pathway; FXS: FC 1.76, p=0.0005; RTT: FC 1.78, p=0.001; DS: FC 2.37, p<0.0001) and Apoe (FXS: FC 1.75, p=0.003; RTT: FC 1.68, p=0.001; DS: FC 2.82, p<0.0001), from all 3 ND astrocytes compared to WT.

Having determined that IP astrocytes express many of the same genes as developing astrocytes in vivo, we then asked if they produce known astrocyte-secreted synapse regulating proteins (Figure 2e-i). Synaptogenic proteins are present at varying abundance in WT ACM. Sparcl1 (silent synapse formation) is most abundant at 0.25%, Gpc4 (immature synapses) is at 0.11%, and Thbs2 (silent synapse formation) is least abundant at 0.002% (Figure 2f)^34^. This demonstrates that protein abundance is not necessarily indicative of functional effect. Comparing protein levels between WT and each ND identified a significant increase in Chrdl1 (synapse maturation) in DS ACM (FC 2.13, p<0.0001)^43^, an increase in Gpc4 and Gpc6 in RTT (Gpc4 FC 1.54, p=0.003; Gpc6 FC 1.65, p=0.001) and DS ACM (Gpc4 FC 1.80, p=0.0008; Gpc6 FC 2.22, p<0.0001), and an increase in Sparcl1 in RTT ACM (FC 1.51, p<0.0001). Only Chrdl1 is also significantly altered at the mRNA level (DS LFC 0.85, p=0.004) (Figure 2g). Conversely Thbs2 mRNA is significantly decreased in FXS astrocytes (LFC −0.81, p=0.04), with no alteration in protein secretion (although Thbs2 protein is low so a change may be below the threshold). For proteins implicated in developmental synapse elimination a similar pattern is seen, with more differences detected in protein level than gene expression (Figure 2h-i). We detected an increase in the extracellular portion of the membrane protein Megf10 (involved in phagocytosis of synapses) in ACM from all NDs (FXS FC 1.57, p=0.005; RTT FC 1.65, p=0.001; DS FC 1.75, p=<0.0001), with no corresponding change in mRNA level. Sparc is already highly abundant in WT ACM (1.95%), and is further upregulated in DS to 3.53% (p<0.00001). Interestingly, two components of the complement cascade that are expressed by astrocytes show opposite changes in DS at both the protein and mRNA level, with C3 increased (protein FC 2.67, p=0.002; mRNA LFC 1.44, p=0.02) and C4b decreased (protein FC −2.02, p=0.0002; mRNA LFC −2.04, p=0.03)^44, 45^.

### Identification of altered protein secretion and gene expression between astrocytes from WT and NDs

The above data demonstrate that IP astrocytes maintain many features of in vivo astrocytes, validating them as a system to identify altered astrocyte protein secretion and gene expression in NDs. We next asked how astrocytes from each individual ND are altered from WT by determining significantly upregulated and downregulated secreted proteins and genes, as well as the overlap between all NDs and WT for each category (Tables S2, S3, S5, S6, S7).

In FXS we found 131 proteins show increased levels in ACM compared to WT, while 547 genes show increased mRNA. Only 4 factors (3.1%) are upregulated at both protein and mRNA levels, including BMP6 (regulates astrocyte maturation; protein FC 2.63, p=0.02; mRNA LFC 0.77, p=0.001) and Sema3e (secreted semaphorin that inhibits neurite outgrowth; protein FC 4.30, p=0.02; mRNA LFC 0.72, p=0.04) (Figure 3a,b)^25, 46^. 108 proteins show decreased level in FXS ACM, and 246 genes are decreased, with only 4 targets (3.7%) overlapping (Figure S3a,b). We found a significant decrease in Fmr1 mRNA in the FXS astrocytes, as would be predicted (LFC - 3.13, p<0.0001). Ntf3 (neurotrophin 3) has been reported to be increased in secretion from FXS astrocytes and to inhibit neurite outgrowth, however we could not detect Ntf3 protein in the ACM^47^. Ntf3 mRNA is present in astrocytes at low levels, with a trend towards an increase in FXS (WT FPKM 0.22, FXS FPKM 0.46, p=0.09). Pathway analysis of the proteins increased in release from FXS astrocytes identified the Alzheimer disease-amyloid secretase pathway, blood coagulation pathway, and regulation of IGF transport and uptake by Igfbps (Figure 3k).

**Figure 3.**
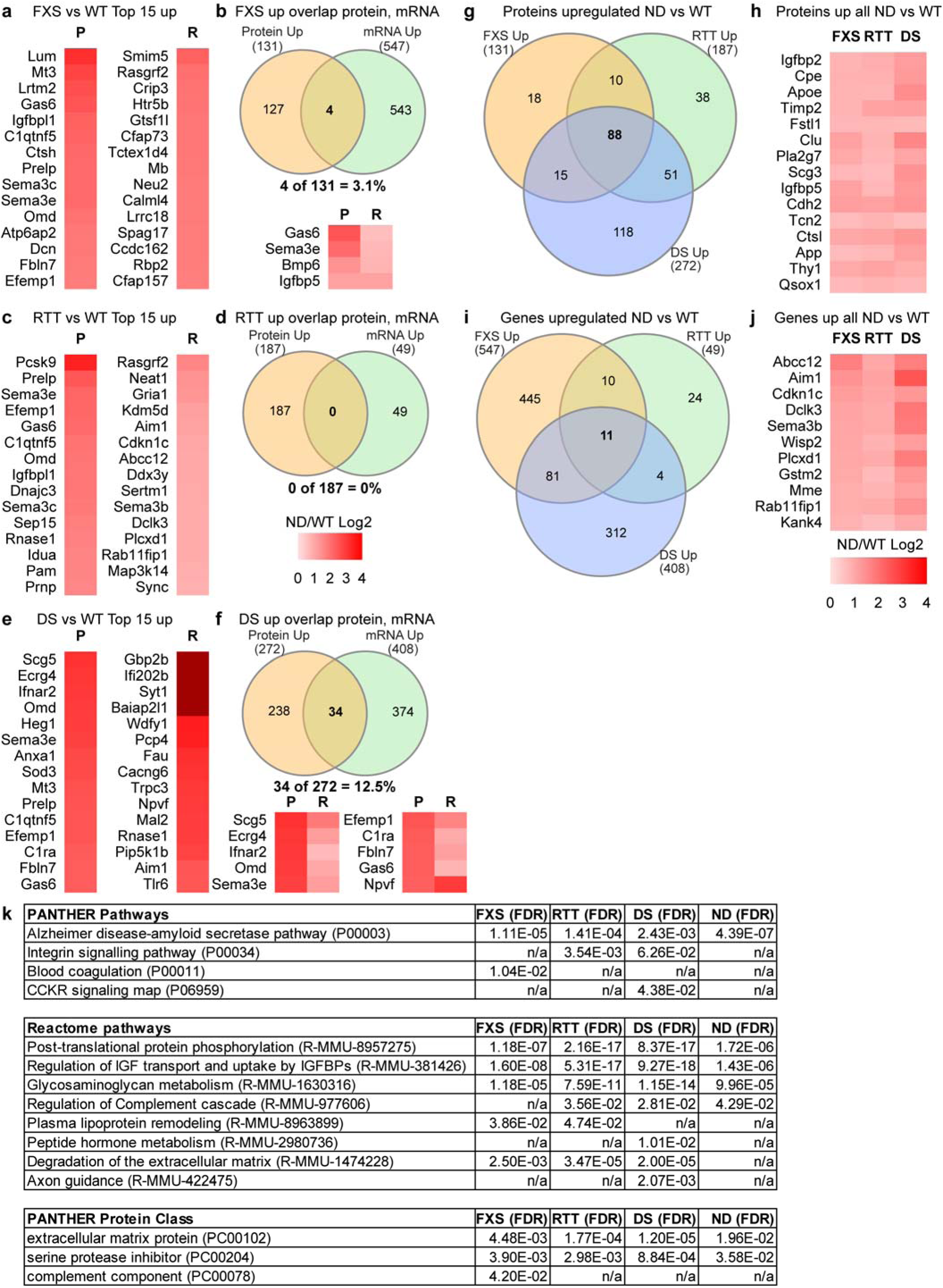
Identification of altered protein secretion and gene expression between astrocytes from WT and NDs. **a,c,e.** Heatmaps of top fold-change in proteins (P) and genes (R) increased between WT and FXS (**a**), RTT (**c**) and DS (**e**) ACM and astrocytes. **b,d,f.** Venn diagram of overlap between proteins and genes with increased level in FXS (**b**), RTT (**d**) and DS (**f**) ACM and astrocytes. **g,h.** Venn diagram showing overlap in proteins increased in all ND (**g**), and heatmap of most abundant altered proteins ranked by protein abundance in FXS ACM (**h**). **i,j.** Venn diagram showing number of genes increased in all ND (**i**) and corresponding heatmap (**j**) of altered genes. **k.** Pathway analysis of proteins upregulated in ND ACM compared to WT demonstrates overlapping alterations in ND astrocyte function compared to WT. Scale bar in **d** applies to all heatmaps, darker red indicates a change above the top of the scale. Proteomics, N=6 cultures per genotype, *p<0.05, abundance >0.01%, fold change between WT and ND ≥1.5. RNASeq, N=6 cultures WT, RTT, FXS; 4 cultures DS, * adjusted p<0.05, FPKM>1, fold change between ND and WT ≥1.5. See also Figure S3; Table S2, S3, S5, S6, S7.

In RTT 187 proteins show increased levels in ACM compared to WT, while only 49 genes show an increase in expression. Interestingly, there is no overlap between the mRNA and protein level increases (Figure 3c,d). Conversely, 200 proteins show a decrease in secretion, while 29 genes have a decrease in mRNA expression (including Mecp2: LFC −5.30, p<0.0001), with an overlap of 1 target (Ktcd12: protein FC −1.68, p=0.03; mRNA LFC −0.61, p=0.007) (Figure S3c,d). Pathway analysis of proteins increased in release from RTT astrocytes identified the Alzheimer disease-amyloid secretase pathway, integrin signaling pathway, and regulation of IGF transport and uptake by Igfbps (Figure 3k).

In DS there are 272 proteins that show increased levels in ACM compared to WT, while 408 genes show increased mRNA, with an overlap of 34 targets (12.5%) (Figure 3e,f). As with FXS, the overlap includes Sema3e (protein FC 7.17, p=0.006; mRNA LFC 1.19, p=0.02) and BMP6 (protein FC 4.07, p=0.0005; mRNA LFC 0.86, p=0.005). 500 proteins show a decrease in DS ACM, while 249 genes are decreased, with 11 targets (2.2%) overlapping (Figure S3e,f). Several genes known to be associated with DS have increased mRNA expression, including App (LFC 1.42, p<0.0001) and Adamts1 (LFC 1.30, p=0.0002)^48^. Additionally, App is increased at the protein secretion level (FC 2.17, p<0.0001). Increased levels of Mt3 and Efemp1 proteins have been reported in astrocytes in DS patients, and we detect increased levels of both in DS ACM (Mt3 FC 5.96, p<0.0001; Efemp1 FC 5.62, p=0.0001)^42, 49^. Pathway analysis of proteins increased in DS ACM identified the amyloid secretase pathway in Alzheimer’s disease, integrin signaling pathway, and regulation of IGF transport and uptake by Igfbps (Figure 3k).

We next asked whether there are proteins or genes that show common changes in all three NDs. We found 88 proteins that are increased in secretion in all 3 NDs vs. WT, while only 11 genes are increased in all 3 NDs, with no overlap in protein and mRNA changes (Figure 3g-j). In contrast, only 32 proteins show a decrease in secretion in all 3 NDs compared to WT, with only 1 gene decreasing, again with no overlap in protein and mRNA (Figure S3g-j). Analysis of the sub-cellular localization of the up and down-regulated proteins by Uniprot determined that 95% of the upregulated proteins are found outside of the cell (dense core vesicle, lysosome, exosome, plasma membrane) whereas only 41% of the downregulated proteins are annotated as extracellular (Table S2). Pathway analysis of upregulated proteins identified the Alzheimer disease-amyloid secretase pathway, regulation of IGF transport and uptake by Igfbps, and regulation of the complement cascade (Figure 3k). Pathway analysis of the gene expression changes found a decrease in both cadherin signaling and Wnt signaling in FXS and RTT astrocytes (Table S6).

The mutations present in RTT, FXS, and DS are all associated with increased expression of genes and/or proteins. Mecp2 (RTT) is a transcriptional repressor, Fmr1 (FXS) is a translational repressor, and in DS there is trisomy of genes^3, 5, 7^. Due to this, combined with results in the literature suggesting ND astrocytes are producing factors that inhibit neurite development^13^, we focused on proteins that show increased secretion in all three NDs for functional testing. Candidate factors that are increased at the protein level in all 3 NDs and have the potential to regulate neurite outgrowth and synapses include Igfbp2, Cpe, App, BMP6 and Sema3e. We chose to focus on Igfbp2, Cpe and BMP6 for further in-depth analysis, to ask if they could inhibit neurite outgrowth in WT neurons. We asked if excess Igfbp2 from astrocytes acts directly on neurons to inhibit neurite outgrowth, due to the positive effect of IGF signaling on neurite outgrowth in RTT and FXS^50, 51^. Cpe is involved in regulated secretion and peptide maturation, including activation of BDNF^52^. Neurons in adult mice lacking Cpe show more complex dendritic arbors compared to WT, so we asked if excess Cpe in ACM is inhibiting neurite outgrowth^53^. As BMP signaling induces structural maturation of astrocytes^25^, we asked if excess BMP6 is acting on astrocytes themselves and is upstream of altered astrocyte protein secretion leading to inhibition of neurite outgrowth. Thus, these candidates represent examples of astrocyte-secreted proteins altered in NDs that act on neurons (Igfbp2, Cpe) and astrocytes (BMP6).

### Excess Igfbp2 in ACM inhibits neurite outgrowth

Our mass spectrometry analysis identified Igfbp2 as a highly abundant protein in WT ACM, that is further significantly increased in release from astrocytes in all 3 NDs (% of total ACM: WT 2.67%, FXS 4.60%, RTT 4.99%, DS 6.16%; Figure 4b,c). Igfbp2 is enriched in astrocytes compared to other cell types in the developing brain (Figure 4d)^54^. Igfbp2 acts in the extracellular space where it binds to IGF1&2 and inhibits their action by preventing them from interacting with the IGF receptor (Figure 4a)^55, 56^. Activation of the IGF pathway in neurons has positive effects, including promoting cellular survival and neurite outgrowth (Figure 4a)^57^, and as summarized in the Introduction, treatment with IGF1 can decrease the neuronal deficits in both RTT and FXS^22–24^. Based on this we hypothesized that increased release of Igfbp2 from ND astrocytes is inhibiting IGF signaling in neurons and contributing to stunted neurite outgrowth.

**Figure 4.**
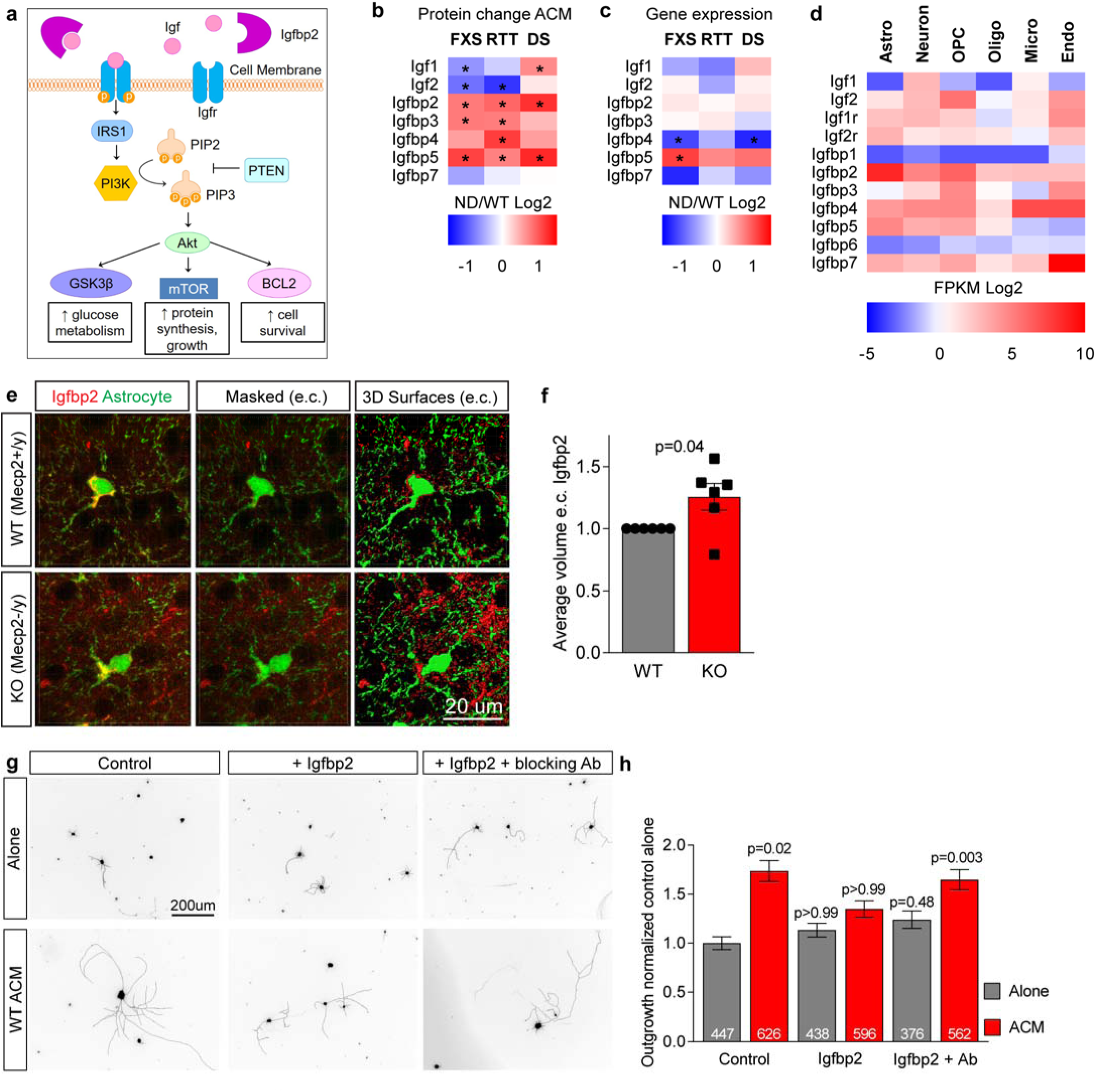
Excess Igfbp2 in ACM inhibits neurite outgrowth. **a.** Schematic of IGF signaling via the PI3K/Akt pathway. **b,c.** Protein secretion (**b**) and gene expression (**c**) profiles of IGF family members in WT and ND astrocytes. Proteomics, N=6 cultures per genotype, *p<0.05, abundance >0.01%, fold change between WT and ND ≥1.5. RNASeq, N=6 cultures WT, RTT, FXS; 4 cultures DS, * adjusted p<0.05, FPKM>1, fold change between ND and WT ≥1.5. **d.** Expression of IGF family members in cortical cell types (data from Zhang et al., 2014). **e,f.** Immunostaining for Igfbp2 in RTT (Mecp2 KO) and WT P7 cortex reveals an increase in extracellular Igfbp2. **e.** Example images (Imaris screenshots) of L2/3 astrocytes (green, Aldh1l1-GFP) immunostained for Igfbp2 (red). Left, confocal image projection; middle, image masked to remove intracellular Igfbp2; right, rendering of Igfbp2 in the extracellular space (e.c.). **f.** Quantification of extracellular Igfbp2. N=6 WT, 6 RTT mice. Bar graph mean±s.e.m. individual data points mice; statistics by T-test. **g,h.** Excess Igfbp2 in ACM inhibits WT neurite outgrowth. **g.** Example images WT neurons grown for 48 hours, conditions as marked. **h.** Quantification total neurite outgrowth normalized to control alone condition. Bar graph mean±s.e.m. Number inside bar = number of neurons, pooled from 3 separate experiments; statistics by one-way ANOVA on ranks, p values against control alone. See also **Figure S4**.

We first asked if Igfbp2 protein is increased in the extracellular space in vivo in mouse models of ND, which would suggest an increase in secretion occurs in vivo as well as in vitro. We focused on Mecp2 KO mice for this analysis, due to the strong links between aberrant IGF signaling and RTT. To do this we crossed Mecp2+/-female mice with male mice expressing Aldh1L1-GFP to mark astrocytes, allowing the identification of astrocytes in brain sections. We compared male littermate Mecp2+/y;Aldh1l1-GFP (WT) and Mecp2-/y;Aldh1l1-GFP (KO) mice at P7, matching the age where cortical astrocytes and neurons are isolated for analysis in vitro. We performed immunostaining for Igfbp2 and used confocal imaging to visualize Igfbp2 and astrocytes in layer (L)2/3 of the visual cortex. We separately analyzed Igfbp2 present within (intracellular, colocalized with GFP) and surrounding (extracellular, no overlap with GFP) astrocytes. This revealed an increase in Igfbp2 levels in the extracellular space (1.26 ± 0.11-fold in RTT vs WT, p=0.04), with no significant change in the volume of Igfbp2 within astrocytes (1.19 ± 0.18-fold in RTT vs WT, p=0.34; Figure 4e,f; S4a). This demonstrates that Igfbp2 is increased in the extracellular space in vivo in RTT, likely due to increased release from astrocytes.

We next asked if increasing the levels of the candidate neurite inhibitory factors Igfbp2 or Cpe in WT ACM is sufficient to inhibit WT neurite outgrowth, by adding excess recombinant Igfbp2 or Cpe protein to WT ACM. For each protein the level in WT ACM was estimated from the mass spectrometry analysis and Western blotting, and recombinant protein added in excess at 4-fold the WT level (Igfbp2 = 240ng/ml; Cpe = 160ng/ml). Treating neurons with Igfbp2 in the absence of ACM had no effect on neurite outgrowth (Alone 253 ± 54µm, Alone + Igfbp2 344 ± 68µm), whereas adding Igfbp2 to WT ACM blocked the neurite-outgrowth promoting effect of WT ACM (WT ACM 661 ± 101µm, WT ACM + Igfbp2 399 ± 64µm; Figure S4b,c). Adding Cpe to WT ACM had no effect on neurite outgrowth, although CPE by itself did promote neurite outgrowth (Alone + Cpe 552 ± 125µm, WT ACM + Cpe 603 ± 100µm; Fig S4b,c). This identifies increased levels of Igfbp2, but not Cpe, in ACM as a potential mediator of inhibited neurite outgrowth in NDs, leading us to focus on Igfbp2.

To ask if excess Igfbp2 in ACM is inhibiting neurite outgrowth by sequestering IGF and decreasing IGF signaling in neurons, we treated WT neurons with WT ACM+Igfbp2+IGF1 (IGF1 = 100 ng/ml). The addition of IGF1 was sufficient to overcome the inhibitory effect of adding excess Igfbp2 to WT ACM, suggesting that Igfbp2 is inhibiting neuronal outgrowth by preventing IGF1 from signaling to neurons (WT ACM + Igfbp2 + IGF1 773 ± 117µm; Figure S4b,c). To further test this hypothesis, and to remove the possibility of the IGF1 rescue being unrelated to Igfbp2, we used an anti-Igfbp2 neutralizing antibody (Igfbp2-Ab) that prevents Igfbp2 from binding to IGF. This was added at 7ug/ml, 2-fold higher than the ND50. WT ACM+Igfbp2 along with the Igfbp2-Ab no longer inhibited neurite outgrowth, providing further evidence that the inhibitory effect of Igfbp2 is through its actions on IGF signaling (total outgrowth normalized to Alone: Alone 1.00 ± 0.07, Alone + Igfbp2 1.13 ± 0.07, Alone + Igfbp2 + Igfbp2-Ab 1.24 ± 0.09, WT ACM 1.74 ± 0.11, WT ACM + Igfbp2 1.35 ± 0.08, WT ACM + Igfbp2 + Igfbp2 Ab 1.65 ± 0.10; Figure 4g,h; S4d,e). A control non-targeting antibody had no effect on WT neurite outgrowth (Figure S4f). This demonstrates the functionality of the Igfbp2-Ab in overcoming the neurite outgrowth inhibiting effects of excess Igfbp2 in ACM. These findings suggest over-production of Igfbp2 by astrocytes is causal to neurite outgrowth deficits in NDs, which we tested next.

### Blocking Igfbp2 in RTT ACM reduces neurite outgrowth deficits

To determine if increased levels of Igfbp2 in ND ACM are contributing to the inhibitory effect on neurite outgrowth, we asked if adding the Igfbp2 blocking Ab to each ND ACM would rescue the outgrowth deficit induced in WT neurons. WT neurons treated with RTT ACM did not show a significant increase in neurite outgrowth compared to neurons alone, whereas the addition of the Igfbp2-Ab to RTT ACM induced a significant increase in neurite outgrowth that partially rescued the deficit (total outgrowth normalized to Alone: Alone 1.00 ± 0.15, WT ACM 3.66 ± 0.38, RTT ACM 1.31 ± 0.16, RTT ACM + Igfbp2-Ab 2.24 ± 0.26; Figure 5a,b; S5a,b). WT neurons cultured with FXS ACM did not show a significant increase in neurite outgrowth compared to neurons alone, demonstrating that FXS ACM is also inhibitory to neurite outgrowth. However, addition of the Igfbp2-Ab to FXS ACM did not significantly enhance neurite outgrowth (total outgrowth normalized to Alone: Alone 1.00 ± 0.07, WT ACM 2.11 ± 0.13, FXS ACM 1.58 ± 0.09, FXS ACM + Igfbp2-Ab 1.69 ± 0.10; Figure 5c,d; S5c,d). We found that DS ACM supports WT neurite outgrowth to a comparable level as WT ACM, and that addition of the Igfbp2-Ab to DS ACM had no effect on neurite outgrowth (total outgrowth normalized to Alone: Alone 1.00 ± 0.06, WT ACM 1.60 ± 0.08, DS ACM 1.53 ± 0.08, DS ACM + Igfbp2-Ab 1.43 ± 0.07; Figure 5e,f; S5e,f). These results show that the inhibitory effect of ND ACM on WT neurite outgrowth is variable, being most severe for RTT ACM, partially inhibitory for FXS ACM, and no inhibition detected with DS ACM (although the reported phenotype in DS is mild, a 15% decrease^58^). We further found that the Igfbp2-Ab could partially rescue the severe inhibition of neurite outgrowth induced by RTT ACM. Of note there is a significant increase in IGF1 secretion from DS astrocytes, which may compensate for the increased Igfbp2 level. Taken together these results demonstrate that in RTT, increased release of Igfbp2 from astrocytes is contributing to the neurite outgrowth deficits.

**Figure 5.**
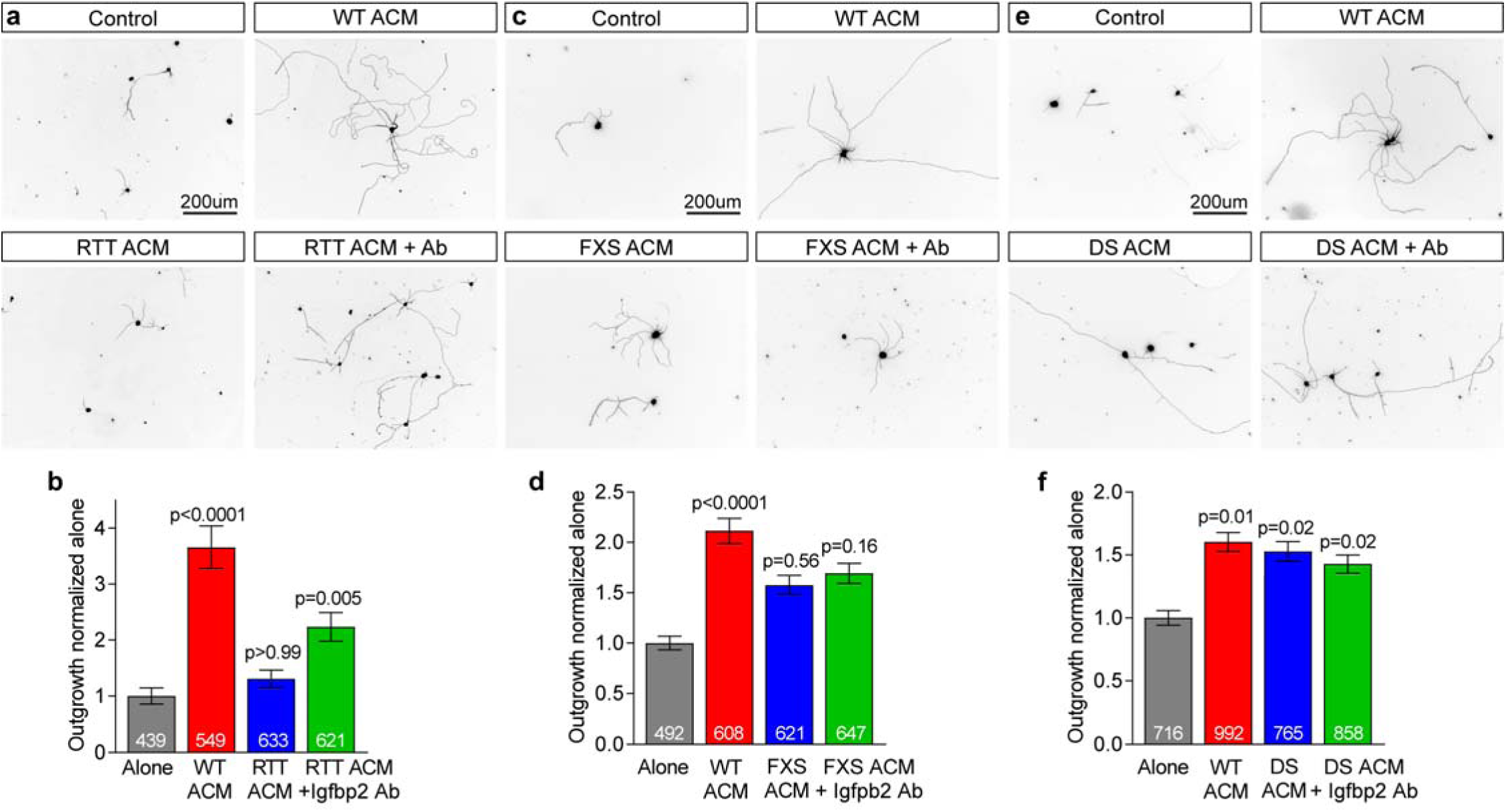
Blocking Igfbp2 in RTT ACM reduces neurite outgrowth deficits. **a-f.** Application of an Igfbp2-neutralizing antibody reduces WT neurite outgrowth inhibition induced by RTT ACM. **a,c,e.** Example images of neurons cultured for 48 hours in RTT (**a**), FXS (**c**) or DS (**e**) ACM. **b,d,f.** Quantification of neurite outgrowth, normalized to alone condition. Bar graphs mean±s.e.m. Number inside bar = number of neurons, pooled from 3 (b), 4 (d), 5 (f) separate experiments. Statistics by one-way ANOVA on ranks, p values against alone. See also **Figure S5**.

### Activating BMP signaling in astrocytes induces changes that overlap with ND astrocytes

Given the partial rescue of WT neurite outgrowth by blocking Igfbp2 in RTT ACM, and the lack of effect of blocking Igfbp2 in FXS ACM, this led us to hypothesize that the combined altered release of multiple proteins from ND astrocytes is responsible for fully inhibiting neurite outgrowth. We reasoned that to target multiple proteins simultaneously we should ask what upstream pathways are commonly altered in ND astrocytes. We identified that BMP6 protein is increased in ACM from all 3 NDs vs WT (Figure 6b). BMP signaling in astrocytes is sufficient to induce morphological maturation of the cells, so we focused on BMP signaling in ND astrocytes themselves^25^. We detected a significant increase in expression of BMP target genes including Id3, Id4, and Smad9 in astrocytes from FXS (Id3: LFC 0.75, p=0.002; Id4: LFC 0.75, p=0.009; Smad9: LFC 0.69, p=0.001) and DS mice (Id3: LFC 1.22, p<0.0001; Id4: LFC 0.95, p=0.002; Smad9: LFC 1.13, p<0.0001), with a trend towards increased Id3 and Smad9 in RTT astrocytes (Figure 6c) (Id3: LFC 0.57, p=0.08; Smad9: LFC 0.50, p=0.1). As canonical BMP signaling regulates expression of many genes, we hypothesized that enhanced BMP signaling in ND astrocytes is upstream of protein secretion changes in these disorders.

**Figure 6.**
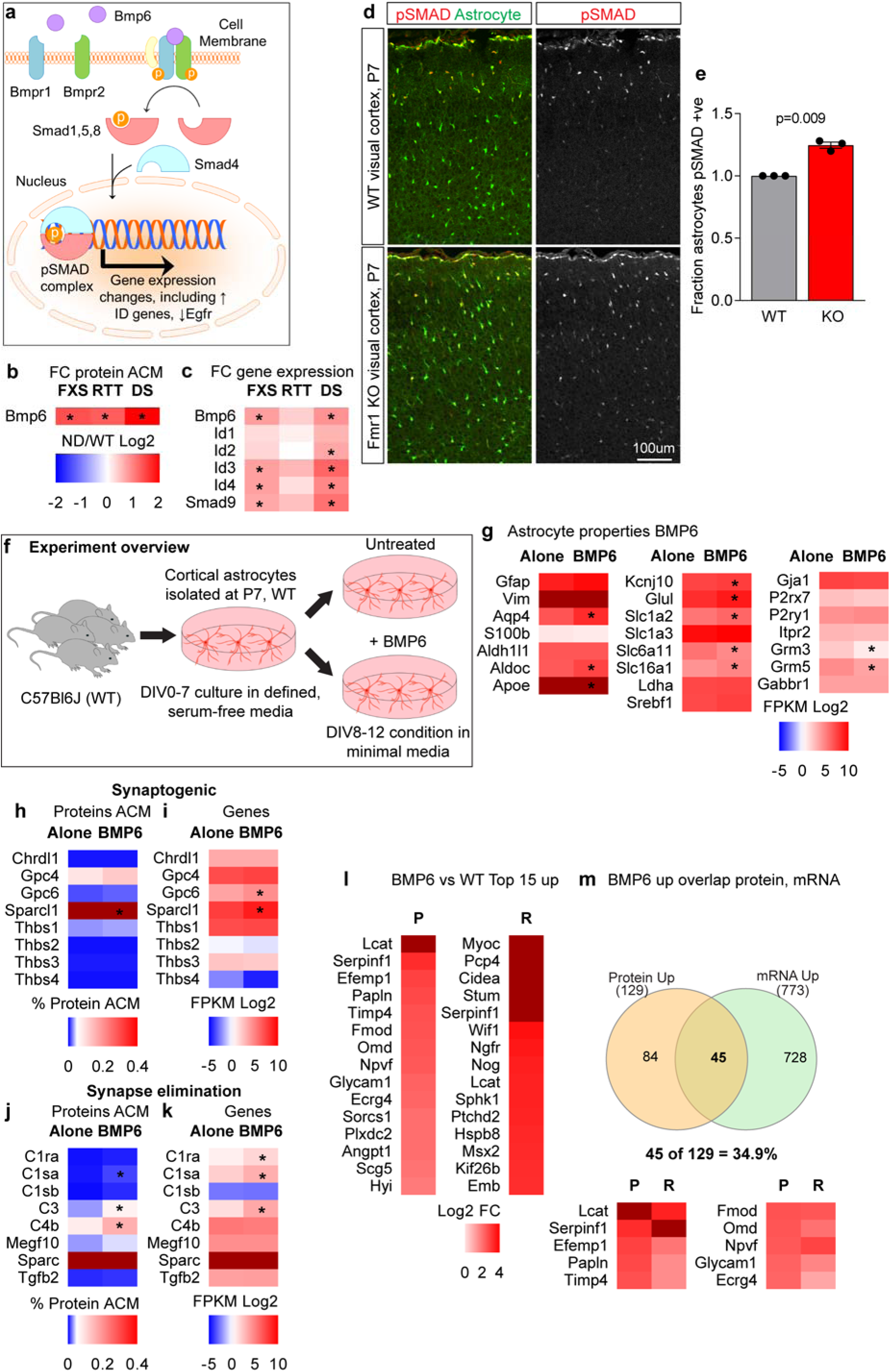
Activating BMP signaling in astrocytes induces changes that overlap with ND astrocytes. **a.** Schematic of canonical BMP signaling pathway. **b.** Fold change in BMP6 protein in ND ACM compared to WT. **c.** Fold change in gene expression for BMP family members in ND astrocytes compared to WT. Proteomics, N=6 cultures per genotype, *p<0.05, abundance >0.01%, fold change between WT and ND ≥1.5. RNASeq, N=6 cultures WT, RTT, FXS; 4 cultures DS, * adjusted p<0.05, FPKM>1, fold change between ND and WT ≥1.5. **d,e.** Increase in the proportion of pSMAD+ astrocytes in FXS visual cortex at P7. **d.** Example images of astrocytes (green, Aldh1l1-GFP) and pSMAD (red) in the visual cortex at P7 of WT (Fmr1+/y) and KO (Fmr1-/y). **e.** Quantification of the proportion of astrocytes that are positive for pSMAD. N=3 WT, 3 FXS mice; bar graph mean±s.e.m., individual data points mice; statistics by T-test. **f.** Experimental schematic for BMP6 treatment of WT astrocytes. **g-m.** Characterization of protein secretion and gene expression profiles of BMP6-treated WT astrocytes compared to untreated WT. **g.** Heatmap of astrocyte identity and function markers. **h-k.** Heatmaps of synaptogenic proteins (**h**) and genes (**i**), as well as synapse eliminating proteins (**j**) and genes (**k**). **l.** Heatmaps of top proteins (P) and mRNA (R) increased between BMP6-treated and untreated WT ACM and astrocytes, ranked by fold-change. For the heatmaps a darker shade of red indicates a value above the top of the scale. **m.** Venn diagram showing overlap between proteins and genes with increased expression in WT astrocytes following BMP6 treatment. For mass spectrometry and RNA Sequencing N=6 cultures, half of each culture treated with BMP6 and other half left untreated. Proteomics, *p<0.05, abundance >0.01%, fold change ≥1.5. RNASeq, * adjusted p<0.05, FPKM>1, fold change ≥1.5. See also Figure S6; Tables S1-7.

To determine if increased expression of BMP target genes is also occurring in ND astrocytes in vivo, we examined expression of phosphorylated SMAD 1/5/8 (pSMAD) in astrocytes in the visual cortex of Fmr1 KO and WT littermates at P7 (FXS). pSMAD is a downstream target of BMPs, and an increase in canonical BMP signaling is reflected by increased pSMAD translocation to the nucleus (Figure 6a)^59^. To do this we crossed Fmr1+/-female mice with male mice expressing Aldh1L1-GFP to mark astrocytes, allowing the identification of astrocytes in brain sections. We compared male littermate Fmr1+/y;Aldh1l1-GFP (WT) and Fmr1-/y;Aldh1l1-GFP (KO) mice at P7, to match the age of the cells used for in vitro analysis. We performed immunostaining for pSMAD and used confocal imaging to visualize pSMAD and astrocytes in the visual cortex. We analyzed the total number of astrocytes and the number of astrocytes positive for pSMAD, and this identified a significant increase in the fraction of astrocytes positive for pSMAD in the cortex in FXS (1.25 ± 0.02-fold in FXS vs WT, p=0.009; Figure 6d,e). These results demonstrate increased BMP signaling is occurring in vivo in astrocytes in FXS KO mice at P7, supporting the hypothesis that aberrant BMP signaling is involved in this disorder.

Given the increase in BMP6 protein secretion from ND astrocytes, activation of BMP target genes in ND astrocytes, and that astrocytes express all components of the canonical BMP pathway (Figure S6a), this led us to ask whether increased BMP6 is acting on astrocytes themselves to change their properties. We predicted that the activation of canonical BMP signaling is upstream of protein secretion alterations in ND astrocytes. To test this hypothesis we treated WT astrocytes with BMP6 (10ng/ml) during the conditioning period (DIV8-12), and compared them to astrocytes from the same culture that were left untreated (Figure 6f). WT astrocytes treated with BMP6 showed distinct changes in morphology (thinner elongated processes), and expressed higher levels of Aqp4 and GFAP protein compared to untreated astrocytes (Figure S6b), reproducing the published finding that BMP signaling induces structural maturation of astrocytes^25^.

We then used mass spectrometry and RNA sequencing to ask if protein secretion and gene expression of WT astrocytes is altered by treatment with BMP6, comparing WT astrocytes with or without BMP6 from 6 separate cultures (Figure 6f). We used the same criteria as described for ND astrocytes to determine significant alterations in protein secretion or gene expression. BMP6-treated astrocytes express the same astrocyte marker genes as their untreated counterparts, with some significant changes in level of expression. The gene expression of Aqp4 is significantly increased in astrocytes by BMP6 treatment (LFC 2.19, p<0.0001), and other genes associated with astrocyte maturation are also increased, including Apoe (LFC 0.62, p=0.003) and Aldoc (LFC 1.10, p<0.0001). Genes downregulated following BMP6 treatment include metabotropic glutamate receptors Grm3 (LFC −0.94, p=0.006) and Grm5 (LFC −0.96, p=0.0005), demonstrating a change in expression similar to that seen in FXS astrocytes (Figure 6g). BMP6 treatment did not alter the majority of synaptogenic factors produced by astrocytes, except for a significant increase in protein secretion and gene expression of Sparcl1 (protein FC 2.17, p=0.005; mRNA LFC 1.32, p<0.0001) (Figure 6h,i). BMP6-treated astrocytes secrete significantly more complement component proteins associated with synapse elimination, including C1sa, C3, and C4b, with a corresponding increase in gene expression for C1sa and C3 (C1sa protein FC 3.23, p=0.004, mRNA LFC 1.41, p<0.0001; C3 protein FC 2.26, p<0.0001, mRNA LFC 1.97, p<0.0001; C4b protein FC 2.12, p=0.0008, mRNA LFC 0.56, p=0.39) (Figure 6j,k). These results indicate that BMP6-treated astrocytes may disrupt synaptic stability by promoting synapse elimination. Overall, BMP6 treatment induces a significant increase in secretion of 129 proteins, and a significant increase in expression of 773 genes, with 45 (34.9%) of the proteins showing increased secretion also showing increased gene expression (Table S2,5,7) (Figure 6l,m). We found a significant decrease in secretion of 273 proteins and a decrease in 731 genes, with 18 (6.6%) of the decreased proteins also showing decreased gene expression (Figure S6c,d).

### Blocking BMP signaling in FXS astrocytes abolishes neurite outgrowth deficits

Given our hypothesis that increased BMP signaling is upstream to alterations detected in ND astrocytes, we next compared the proteins altered in secretion in all 3 NDs with those altered by BMP6 treatment of WT astrocytes. 38 of the 88 proteins that are increased in ND ACM are also increased in BMP6-ACM (43.2%), whereas 14 of the 32 downregulated proteins are also downregulated by BMP6 (43.8%) (Figure 7a; S7a; Table S2). Furthermore, of the 11 genes that show increased expression in all NDs compared to WT, 4 of them are also increased in BMP6-treated astrocytes, and the 1 gene showing decreased expression in all NDs is also decreased following BMP6 treatment (Figure 7b; S7b). This shows a strong overlap in the protein secretion and gene expression profiles of ND astrocytes and WT astrocytes treated with BMP6, indicating that BMP signaling is upstream of many of the protein release alterations in astrocytes in ND. Interestingly Igfbp2 and Igfbp5 both show increased secretion from BMP6-treated WT astrocytes, matching the change seen in NDs (Igfbp2 FC 1.50, p=0.006; Igfbp5 FC 1.77, p=0.01) (Figure 7c). Pathway analysis of genes upregulated by BMP6 treatment also identified the Igfbp pathway (Table S6), providing further support for BMP6 inducing upstream changes in astrocyte function that contribute to aberrant IGF signaling in NDs.

**Figure 7.**
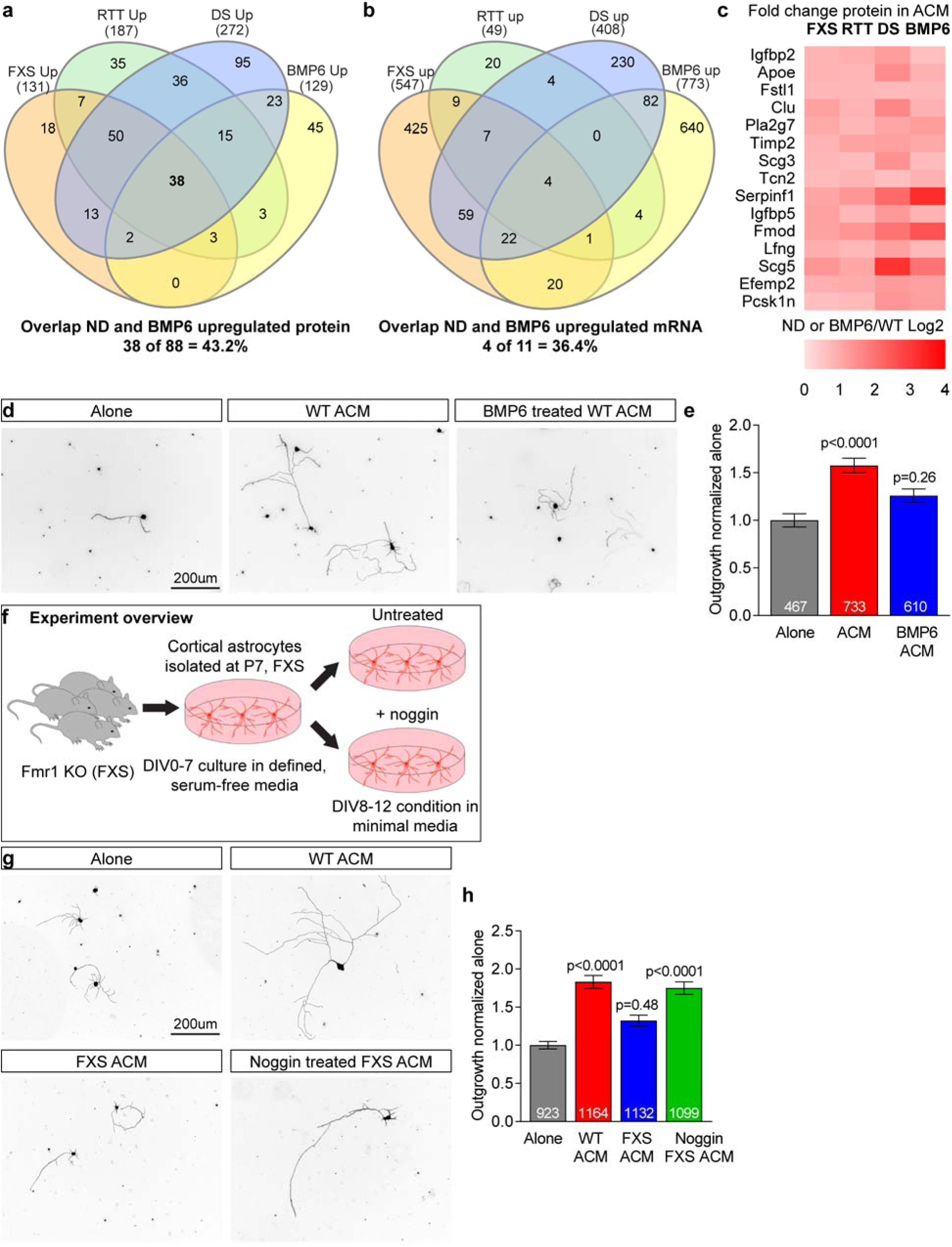
Blocking BMP signaling in FXS astrocytes abolishes neurite outgrowth deficits. **a-c.** BMP6-treated WT astrocytes show increased protein secretion (**a**) and gene expression (**b**) that overlaps with ND astrocytes. **c.** Heatmap of proteins increased in all ND and BMP6-treated astrocytes vs. WT, ranked by protein abundance in BMP6-treated ACM. N=6 cultures each WT, FXS, RTT, DS, plus 6 cultures WT +/-BMP6. **d,e.** ACM from WT astrocytes treated with BMP6 inhibits WT neurite outgrowth. **d.** Example images of WT neurons cultured for 48 hours, conditions as marked. **e.** Quantification of total neurite outgrowth. N = 3 experiments. **f-h.** Blocking BMP signaling in FXS astrocytes rescues deficits in WT neurite outgrowth. **f.** Experimental schematic for noggin treatment of FXS astrocytes. **g.** Example images of WT neurons cultured for 48 hours, conditions as marked. **h.** Quantification of total neurite outgrowth. N=3 experiments. Bar graphs represent mean±s.e.m. Numbers inside bar = number of neurons. Statistics by one-way ANOVA on ranks, p-value compared to alone. See also Figure S7; Table S2, S5.

Given the overlap in altered protein secretion induced in WT astrocytes by BMP6-treatment with those seen in ND ACM, we next asked if ACM from BMP6-treated WT astrocytes is sufficient to inhibit outgrowth of WT cortical neurons. Neurons were cultured for 48 hours alone, with WT ACM or BMP6-ACM, and neurite outgrowth analyzed. This demonstrated that BMP6-ACM is not able to enhance neurite outgrowth compared to neurons grown alone, whereas untreated WT ACM can (total outgrowth normalized to Alone: Alone 1.00 ± 0.07, WT ACM 1.58 ± 0.08, BMP6-ACM 1.26 ± 0.07; Figure 7d,e; S7c,d). This data strongly suggests that aberrant BMP signaling in ND astrocytes is upstream of secretion alterations that lead to changes in astrocyte support of neurite outgrowth and neuronal development.

Based on this we next asked if inhibiting BMP signaling in ND astrocytes would rescue the inhibitory effects of ND ACM on WT neurite outgrowth. We chose to focus on FXS for these experiments, as we identified increased BMP pathway activation in astrocytes in FXS both in vitro and in vivo (Figure 6b-e). We isolated FXS astrocytes and treated half of them during the conditioning period with the secreted BMP inhibitor noggin (1ug/ml) to bind BMPs and prevent BMP-receptor interaction, while leaving the other half untreated (Figure 7f)^59^. ACM from untreated FXS astrocytes did not induce a significant increase in WT neurite outgrowth compared to neurons alone, whereas ACM from FXS astrocytes treated with noggin supported neurite outgrowth to a level indistinguishable from WT ACM (total outgrowth normalized to Alone: Alone 1.00 ± 0.05, WT ACM 1.83 ± 0.08, FXS ACM 1.32 ± 0.07, noggin-FXS ACM 1.75 ± 0.08; Figure 7g,h; S7e,f). To ask if noggin may be having a direct effect on neurons independent of its effect on FXS astrocytes, we added purified recombinant noggin to WT and FXS ACM at the time of treatment of cortical neurons. This showed that noggin added directly to neurons does not enhance neurite outgrowth, indicating that it is the effects of noggin on FXS astrocytes during conditioning that allows noggin-treated FXS ACM to support neurite outgrowth (Figure S7g,h). These results support a critical role for increased BMP signaling in astrocytes in FXS in regulating aberrant protein secretion, that leads to inhibition of neuronal development.

## Discussion

In this work we generated a resource to ask how alterations to astrocyte protein secretion contribute to aberrant neuronal development in diverse NDs. To achieve this we developed an approach to isolate cortical astrocytes and neurons from mice at the end of the first postnatal week, a time point when astrocytes regulate neuronal outgrowth and synapse formation in vivo. We maintained the astrocytes in vitro in defined conditions that maintain their in vivo properties and prevent them from becoming reactive. By combining quantitative mass spectrometry analysis of proteins secreted by astrocytes with RNA sequencing analysis of gene expression, we identified the normal protein secretion and gene expression profile of astrocytes at this timepoint, along with how these are altered in multiple genetic forms of NDs. We identified a number of differences in both protein secretion and gene expression between each individual ND and WT, as well as 88 proteins that are upregulated in their secretion from all 3 NDs. Further testing showed two of these upregulated proteins, Igfbp2 and BMP6, have negative effects on neuronal development. Analysis of RNA sequencing determined pathways altered in ND astrocytes, leading us to identify that enhanced BMP signaling is upstream of many ND secretion alterations, including Igfbp2.

While we identified many alterations in protein release from ND astrocytes, the majority of these changes do not show a corresponding alteration in gene expression. This highlights the importance of a proteomics approach when studying inter-cellular signaling interactions, particularly in the case of astrocytes, which are a cell type specialized for secretion^21, 60^. The proteins we detected in the extracellular fraction come from a number of sources including dense core vesicles, lysosomes, exosomes and cleaved membrane proteins. The proteins that are increased in release from ND astrocytes come from all of these sources, indicating that there is not one particular release pathway that is altered in astrocytes in NDs, rather an overall change to the cell is occurring. As we discuss below, we identified that BMP signaling in ND astrocytes is upstream of nearly half of the protein secretion changes, showing this is one important regulator. The pathways that control altered release of the other 50% of proteins remain to be determined. Our RNA sequencing analysis identified decreased activation of the Wnt pathway in astrocytes in all 3 NDs, providing one example candidate mechanism for future investigation.

We identified that increased extracellular levels of astrocyte-derived Igfbp2 are contributing to stunted neurite outgrowth in RTT. The IGF pathway shows decreased activity in RTT brains^61^, and IGF signaling is altered in multiple different NDs. Studies of patients in a number of NDs, however, have not found consistent alterations in IGF1, with increased, decreased or unchanged levels detected^62, 63^. This suggests factors other than the overall level of IGF1 itself are contributing to the signaling deficit, such as an upregulation in extracellular binding proteins. Transgenic mice overexpressing Igfbp2 have a significant decrease in body weight as well as a modest decrease in brain weight, suggesting Igfbp2 is normally inhibiting IGF signaling and suppressing growth^55^. Further, in Pallister-Killina syndrome, associated with impaired postnatal growth and intellectual impairment, there is evidence for increased Igfbp2 in the serum of some patients^56^. There are six high affinity Igfbps (Igfbp1-6). In addition to Igfbp2, we detected a significant increase in release of Igfbp3, 4 and 5 from RTT astrocytes, and a decrease in IGF2. Increased Igfbp3 has also been reported in the brains of Mecp2 KO mice and RTT patients^64^. In FXS ACM Igfbp3 and 5 are increased, and IGF1 and 2 decreased. In DS ACM there is an increase in Igfbp5 along with increased IGF1. This shows alterations to the IGF-Igfbp pathway are present in astrocytes in all NDs but at variable levels, being most affected in RTT, less so in FXS, and the least affected in DS where there is also a potentially compensatory increase in IGF1. This suggests a general dysregulation of Igfbps is occurring in astrocytes in NDs, which may contribute to the decreased IGF pathway activation that is observed. Developing ways to block the interaction of specific Igfbps with IGF, in order to liberate IGF in the appropriate location for it to signal, may provide a new approach to enhance IGF signaling in multiple disorders.

We demonstrated that increased BMP pathway activity is upstream of much of the altered protein release from ND astrocytes, including Igfbp2, identifying an important role for the canonical BMP pathway in ND astrocytes. Not much is known about the role of BMP signaling in NDs, however Bmpr2 is a target of Fmr1, and there is a partial rescue of spine deficits in FXS mice that are heterozygous for Bmpr2^65^. This spine rescue is thought to be due to decreased non-canonical BMP signaling in FXS neurons, however our data suggest a role for canonical BMP signaling in astrocytes should also be considered. Dysregulated BMP signaling has been implicated in other NDs, for example there is an increase in BMP6 in the serum of patients with Asperger’s Syndrome^66^. Due to these findings, the development of approaches to specifically decrease BMP signaling in astrocytes may be beneficial in multiple NDs.

While we have focused on Igfbp2 and BMP6, there are a number of other proteins altered in ND ACM that may contribute to aberrant neuronal development. Class 3 secreted semaphorins are upregulated in ACM from all 3 NDs, and these are inhibitory to neurite outgrowth^46^. Neuronal Sema3f has previously been linked to both RTT and FXS, providing further rationale for examining the role of astrocyte secreted semaphorins in NDs^67, 68^. Additionally, while we focused on upregulated proteins, we also identified a number of proteins that show a decrease in all three NDs versus WT. These include Sulf2 (sulfatase 2, regulates heparan sulfate proteoglycans), Hdgfrp3 (hepatoma derived growth factor related protein 3, a neurotrophic factor), and Ptn (pleiotrophin). All three of these proteins are known to support neuronal development and promote neurite outgrowth^69–71^, so their deficiency may further contribute to the observed failure of WT neurons to develop in the presence of ND ACM. We also found that decreased release of Hdgfrp3 and Ptn from ND astrocytes is downstream of BMP signaling, further linking upregulated BMP signaling in astrocytes in NDs to aberrant neuronal development.

We found an upregulation of the Alzheimer’s disease-amyloid secretase pathway, and in particular increased secretion of App, Apoe and Clu, from astrocytes in all 3 NDs. App is known to be increased in DS as it is present on the triplicated chromosome^72^. App is also a target of Fmr1, and increased App protein has been observed in Fmr1 KO neurons in vitro^73, 74^. We found that App is also increased at the protein level in RTT. In all three disorders we find astrocytes are upregulating App, providing an alternative source of the protein to neurons. Beyond development, DS is associated with early onset Alzheimer’s Disease^75^, whereas fragile X associated tremor ataxia syndrome (FXTAS) can occur in people with Fmr1 mutations^76^. In addition, Igfbp2 is increased in the cerebrospinal fluid of patients with Alzheimer’s disease, and in astrocytes in mouse models of Alzheimer’s disease^77^. Together this demonstrates that the upregulation of proteins normally associated with neurodegeneration (App, Apoe, Clu, Igfbp2), is also occurring in neurodevelopmental disorders. This suggests that these proteins may represent a general response of astrocytes across neurological disorders, from neurodevelopmental to neurodegenerative.

We chose to analyze astrocytes from FXS, RTT and DS due to the phenotypic overlap in the neuronal alterations in these conditions: a decrease in neurite outgrowth and impaired synaptic development. Our in vitro experiments demonstrated that the effect of astrocytes from these NDs on neurite outgrowth is variable, matching what has been reported for neuronal deficits. RTT ACM has the most severe effect and fails to support any neurite outgrowth; FXS ACM supports some neurite outgrowth but it is less than WT; and DS ACM does not inhibit neurite outgrowth. Our functional analysis of protein secretion changes identified possible mechanisms underlying these differences, for example over-lapping yet distinct alterations to the IGF-Igfbp pathway in ND astrocytes (discussed above). We focused on neurite outgrowth to determine the inhibitory effects of ND ACM, whereas in the case of DS the strongest reported effect is on dendritic spines. Extending the functional analysis to look at spine and synapse development will determine what shared astrocytic alterations in NDs impact later stages of neuronal development. Furthermore, astrocytes are not always inhibitory to neurons in NDs. In Costello Syndrome (CS), a ND caused by mutations in HRAS, mutant astrocytes have the opposite effect and increase neurite outgrowth and synapse formation^78^. CS astrocytes upregulate release of a different set of proteins than we identified, predominantly extracellular matrix factors such as agrin, brevican and neurocan. These protein secretion differences exemplify the importance of using a shared phenotype as the basis for comparative studies of astrocytes across NDs, in order to identify changes that are likely to impact the same pathways in neurons. Indeed, recent work has proposed that common mechanisms may underlie overlapping neuronal phenotypes in DS and FXS^79^, and we propose this comparative approach should also be applied to astrocytes in NDs, as we have done here.

In summary, by combining a physiologically relevant cell culture system with quantitative proteomics analysis of secreted proteins, we have identified how astrocytes alter their interaction with neurons in multiple genetic neurodevelopmental disorders. This analysis has identified candidate non-neuronal proteins and pathways that can be targeted in these disorders, as well as in disorders of unknown cause with shared phenotypes, in order to correct aberrant neuronal development and function.

## Online Methods

Step by step protocols, including for immunopanning isolation of mouse astrocytes and neurons, are available on request from. The immunopanning protocols will also be made available via www.protocols.io

### Animals

All animal work was approved by the Institutional Animal Care and Use Committee (IACUC) of the Salk Institute for Biological Studies.

### Mice

Mice were housed in the Salk Institute animal facility at a light cycle of 12 hr light: 12 hr dark, with access to water and food ad libitum.

#### Wild-type (WT) mice

WT C57BL/6J (Jax 000664) mice were used to generate WT astrocytes in vitro, to generate neurons for all cortical neuronal assays, and to breed to Mecp2 and Fmr1 mice. Mice of both genders were used.

#### Mecp2 knockout (KO) mice

B6.129P2(C)-Mecp2tm1.1Bird/J (Jax 003890) mice were used to generate RTT astrocytes in vitro, to breed Mecp2 mice, and to breed Aldh1L1-eGFPxMecp2 mice^33^. Mecp2 is found on the X chromosome and the hemizygous condition is fatal to juvenile males, so experimental mice were generated by breeding heterozygous (+/-) Mecp2 females to WT (+/y) C57BL/6J males. Astrocytes were isolated from male Mecp2 (-/y) mice.

#### Fmr1 KO mice

B6.129P2-Fmr1tm1Cgr/J (Jax 003025) mice were used to generate FXS astrocytes in vitro, to breed Fmr1 mice, and to breed Aldh1L1-eGFPxFmr1^31^. Fmr1 is found on the X chromosome and KO males are fertile, so experimental mice were generated by breeding heterozygous (+/-) Fmr1 females to KO (-/y) males. Astrocytes were isolated from both male and female KO mice.

#### Down syndrome transgenic mice

B6EiC3Sn.BLiA-Ts(1716)65Dn/DnJ (Jax 005252) mice were used to generate DS astrocytes in vitro, and to breed Ts65Dn mice^32, 80^. Ts65Dn mice are trisomic for two-thirds of the genes orthologous to human chromosome 21 and only one copy of the mutation is required for the condition, so experimental mice were generated by breeding Ts65Dn+ female mice to WT males. Astrocytes were prepared from both male and female Ts65Dn+ mice.

#### Aldh1l1-EGFP x Mecp2 KO or Fmr1 KO mice

Tg(Aldh1l1-EGFP)OFC789Gsat/Mmucd mice express Aldh1l1-eGFP in astrocytes (011015-UCD). Male Aldh1l1-eGFP mice were bred to Mecp2 or Fmr1 heterozygous (+/-) females. Mecp2 and Fmr1 KO male mice and their WT male littermates expressing eGFP in astrocytes were used for experiments.

### Cell culture

#### Cortical astrocyte cultures

Cortical mouse astrocytes were isolated by immunopanning from WT, RTT, FXS, and DS mice, using an adapted version of a protocol developed for rat astrocytes^20, 26^. The cortices of 2-4 mouse pups (P5-P7) were dissected out and the meninges removed, before tissue was digested in a papain enzymatic solution to generate a single cell suspension. This was then passed over a series of five plates to deplete unwanted cell types (lectin [Vector Labs Inc, #L-1100], 10 minutes, to deplete endothelia; cd11b [Ebioscience, #14-01112-86], 10 minutes, to deplete microglia, cd45 [BD Pharmingen, #550539], 20 minutes, to deplete macrophages, and O4 hybridoma [see^81^ for recipe], 15 minutes, x2, to deplete oligodendrocyte precursor cells) before positive selection for astrocytes using the astrocyte cell surface antigen 2 (ACSA2, Miltenyi Biotech #130-099-138). Astrocytes were plated on glass coverslips (12mm diameter, Carolina Biological Supply 633029) coated with poly-D-lysine (Sigma P6407) in 24 well plates (Falcon 353047) at a density of 50,000-80,000 cells/well or in six well plates (Falcon 353046) coated with poly-D-lysine at a density of 280,000 - 350,000 cells/well in a growth medium containing: 50% DMEM (Thermo Fisher Scientific 11960044), 50% Neurobasal (Thermo Fisher Scientific 21103049), Penicillin-Streptomycin (LifeTech 15140-122), GlutaMax (Thermo Fisher Scientific 35050-061), sodium pyruvate (Thermo Fisher Scientific 11360-070), N-acetyl-L-cysteine (Sigma A8199), SATO (containing: transferrin (Sigma T-1147), BSA (Sigma A-4161), progesterone (Sigma P6149), putrescine (Sigma P5780), sodium selenite (Sigma S9133)), and heparin binding EGF like growth factor (HbEGF, R&D Systems 259-HE/CF). Astrocyte cultures were maintained in a humidified incubator at 37°C and 10% CO2 and grown to confluence (5-7 days).

#### Generation of astrocyte conditioned media (ACM)

Astrocytes grown in 6 well plates were washed 3 times with warm DPBS (HyClone SH30264) to remove any minor traces of added protein from growth media, and the growth media was replaced by minimal low protein conditioning media (50% DMEM, 50% Neurobasal, penicillin-streptomycin, glutamax, sodium pyruvate, N-acetyl-Lcysteine (Sigma A8199), and carrier-free HbEGF (R&D Systems, 259-HE/CF, resuspended in dPBS only with no BSA). Astrocytes were conditioned in low protein media for 5 days in a tissue culture incubator at 37°C, 10% CO2. The astrocyte conditioned media (ACM) was collected and concentrated 30-fold using Vivaspin 20 or 6 centrifugal concentrators with a MW cut-off of 3 kDa (Sartorius VS2052 or VS0652). Protein concentration of the ACM was assessed by Bradford protein assay (Bio-Rad 500-0006). ACM for mass spectrometry and Western blotting was flash-frozen with liquid nitrogen and stored at −80°C until analysis. ACM for cortical assays was stored at 4°C until use, for no more than 2 weeks.

### Cortical neuron cultures

Cortical neurons were isolated from P5-P7 C57Bl6J WT mice using immunopanning, modified from^29^. The single cell suspension was passed over two plates to deplete unwanted cell types (lectin, IgG H+L [Jackson Immunoresearch #115-005-167]) before positive selection for cortical neurons using the neuronal marker NCAM-L1 (Millipore MAB5272). Cortical neurons were plated on glass coverslips (12mm diameter, Carolina Biological Supply 633029) coated with poly-D-lysine (Sigma P6407) and laminin (R&D 340001001) at a density of 50,000-80,000 cells/well in a minimal medium, with the addition of astrocyte conditioned media and/or protein factors and antibodies as described in the text. Cortical neuron minimal media contained: 50% DMEM (Thermo Fisher Scientific 11960044), 50% Neurobasal (Thermo Fisher Scientific 21103049), Penicillin-Streptomycin (LifeTech 15140-122), glutamax (Thermo Fisher Scientific 35050-061), sodium pyruvate (Thermo Fisher Scientific 11360-070), N-acetyl-Lcysteine (Sigma A8199), insulin (Sigma I1882), triiodo-thyronine (Sigma T6397), SATO (containing: transferrin (Sigma T-1147), BSA (Sigma A-4161), progesterone (Sigma P6149), putrescine (Sigma P5780), sodium selenite (Sigma S9133)), B27 (see^82^ for recipe), and forskolin (Sigma F6886). Cortical neuron cultures were maintained in a humidified incubator at 37°C and 10% CO2.

### Treatment with protein factors

Unless stated otherwise in the text, all treatments were applied for 48 hours before cortical neurons were analyzed. In each experiment there was a negative control condition, cortical neurons in minimal media (alone condition; treated with buffer), and a positive control condition, cortical neurons treated with WT astrocyte conditioned media (ACM) at 3 ug/mL (WT ACM condition). The candidate proteins identified by mass spectrometry were insulin like growth factor binding protein 2 (Igfbp2) (R&D 797-B2 resuspended at 100 ug/mL in sterile PBS), carboxypeptidase E (CPE) (Abcam #ab169054 resuspended at 100 ug/mL in sterile deionized water), and BMP6 (R&D 507-BP/CF resuspended at 10 ug/mL in sterile dPBS). Igfbp2 and CPE were tested at a concentration determined by their overall levels in WT and ND ACM, at a level of roughly 4X the concentration expected in WT ACM (2X the concentration expected in ND ACM). Igfbp2 was applied at a concentration of 240 ng/mL, while CPE was applied at a concentration of 160 ng/mL. For BMP6 treatments, BMP6 was added to half of the wells of WT astrocytes during the conditioning phase, for a total of 5 days, at a concentration determined by previously published research^25^, 10 ng/mL, before being collected as normal and applied to cortical neurons at the same concentration of 3 ug/mL. Igfbp2-neutralizing antibody (R&D MAB797 at 0.5 mg/mL in sterile PBS) and its control IgG (R&D 6-001-F at 0.5 mg/mL in sterile PBS) were used at 7 ug/mL, a concentration determined to be twice the ND50 for the level of Igfbp2 in solution. IGF1 (R&D 791-MG, at 100 ug/mL in sterile PBS) was used at 100 ng/mL, a concentration previous used to study the effects of IGF1 on neurite outgrowth in RTT^22^. For noggin treatments, noggin (R&D 1967-NG/CF at 200 ug/mL in sterile PBS) was added to half of the wells of FXS astrocytes during the conditioning phase, for a total of 5 days, at 1 ug/mL, before being collected as normal. Due to the high levels of residual noggin in the treated ACM, FXS untreated and FXS+noggin ACM were concentrated to the same volume and a protein concentration of 3 ug/mL determined in the untreated ACM; equal volumes of FXS untreated and FXS+noggin ACM were applied to cortical neurons. To test effects of noggin alone, noggin was added at 1 ug/mL directly to the minimal medium immediately before plating cortical neurons.

### Mass spectrometry

Protein concentration of ACM was determined by Bradford Assay and ACM was distributed into aliquots of 15 ug before being flash-frozen with liquid nitrogen and stored at −80°C until analysis. Samples were thawed and split into 3 equal parts of 5 ug each to produce technical triplicates for each biological replicate. Samples were precipitated by methanol/chloroform. Dried pellets were dissolved in 8 M urea/100 mM TEAB, pH 8.5. Proteins were reduced with 5 mM tris(2-carboxyethyl)phosphine hydrochloride (TCEP, Sigma-Aldrich) and alkylated with 10 mM chloroacetamide (Sigma-Aldrich). Proteins were digested overnight at 37°C in 2 M urea/100 mM TEAB, pH 8.5, with trypsin (Promega). Digestion was quenched with formic acid, 5% final concentration. The digested samples were analyzed on a Fusion Lumos Orbitrap tribrid mass spectrometer (Thermo). The digest was injected directly onto a 30 cm, 75 um ID column packed with BEH 1.7um C18 resin (Waters). Samples were separated at a flow rate of 300 nl/min on a nLC 1000 (Thermo). Buffer A and B were 0.1% formic acid in water and 0.1% formic acid in 90% acetonitrile, respectively. A gradient of 1-25% B over 160 min, an increase to 35% B over 60 min, an increase to 90% B over 10 min and held at 100%B for a final 10 min was used for 240 min total run time. Column was re-equilibrated with 20 ul of buffer A prior to the injection of sample. Peptides were eluted directly from the tip of the column and nanosprayed directly into the mass spectrometer by application of 2.5 kV voltage at the back of the column. The Orbitrap Fusion was operated in a data dependent mode. Full MS scans were collected in the Orbitrap at 120K resolution with a mass range of 400 to 1500 m/z and an AGC target of 4e5. The cycle time was set to 3 sec, and within this 3 sec the most abundant ions per scan were selected for CID MS/MS in the iontrap with an AGC target of 1e4 and minimum intensity of 5000. Maximum fill times were set to 50 ms and 200 ms for MS and MS/MS scans respectively. Quadrupole isolation at 1.6 m/z was used, monoisotopic precursor selection was enabled and dynamic exclusion was used with exclusion duration of 5 sec. Protein and peptide identification were done with Integrated Proteomics Pipeline – IP2 (Integrated Proteomics Applications). Tandem mass spectra were extracted from raw files using RawConverter^83^ and searched with ProLuCID^83^ against Uniprot mouse database. The search space included all fully-tryptic and half-tryptic peptide candidates, carbamidomethylation on cysteine was considered as a static modification.

The validity of the peptide spectrum matches (PSMs) generated by ProLuCID was assessed using Search Engine Processor (SEPro) module from PatternLab for Proteomics platform^84^. XCorr, DeltaCN, DeltaMass, ZScore, number of peaks matched, and secondary rank values were used to generate a Bayesian discriminating function. A cutoff score was established to accept a false discovery rate (FDR) of 1% based on the number of decoys. A minimum sequence length of six residues per peptide was required and results were post-processed to only accept PSMs with < 10ppm precursor mass error. Volcano plots were generated by a pairwise comparison of each individual ND versus WT using PatternLab’s TFold module. The following parameters were used to select differentially expressed proteins: spectral count data were normalized using NSAF values^85^, and two nonzero replicate values were required for each condition. A BH q-value was set at 0.05 (5 % FDR). A variable fold-change cutoff for each individual protein was calculated according to the t-test p-value using an F-Stringency value automatically optimized using the TFold software.

### qRT-PCR & RNA sequencing

For RT-PCR, following ACM collection RNA was collected using the RNEasy Micro Plus kit (Qiagen 74034). Aliquots (0.1 - 0.5 ug) of RNA were used for cDNA synthesis by RT-PCR using Superscript VILO master mix (Thermo Fisher Scientific 11755050) according to manufacturer’s instructions. For each sample, an equal amount of whole brain RNA from an animal with a matching genotype (WT, MeCP2 KO, Fmr1 KO, or Ts65Dn Mut) was used to generate control cDNA. 1 uL of the obtained cDNA was used for qPCR reaction using SYBR green PCR mastermix (Life Supply 4309155). All samples were run in triplicates (technical replicates). Primer pairs were as follows: GAPDH (loading control; forward TGCCACTCAGAAGACTGTGG, reverse GCATGTCAGATCCACAATGG); Syt1 (neurons; forward CTGCATCACAACACTACTAGC, reverse CCAACATTTCTACGAGACACAG); GFAP (astrocytes; forward AGAAAACCGCATCACCATTC, reverse TTGAGAGGTCTTGTGACTTTT); CSPG4 (oligodendrocyte precursor cells; forward CTCAGAACCCTATCTTCACG, reverse TACATGGTAGTGGACCTCAT); CD68 (microglia; forward ATACAATGTGTCCTTCCCAC, reverse CTATGCTTGCATTTCCACAG); FGFr4 (fibroblasts; forward AGAGTGACGTGTGGTCTTT, reverse ACTCCCTCATTAGCCCATAC). Only samples that had GFAP expression of ≥3x whole brain and CSPG4 expression of ≤2x whole brain were used for proteomics and gene expression analysis.

For RNA sequencing, following ACM collection RNA was collected using the RNEasy Micro Plus kit (Qiagen 74034). For RNA sequencing RNA libraries were prepared with TrueSeq Stranded mRNA Library Prep Kit (Illumina), from on average 0.8ug total RNA. Poly(A) mRNA was isolated from total RNA samples, followed by mRNA fragmentation, first strand cDNA synthesis, second cDNA synthesis and adaptor ligation, and isolation and amplification of cDNA fragments. Illumina HiSeq 2500 instrument was used for sequencing, with a read length of 50bp single-end, and an average of 36 million reads per sample.

Sequenced reads were quality-tested using FASTQC v0.11.8 and aligned to the mm10 mouse genome using the STAR aligner version 2.5.3a. Mapping was carried out using default parameters, filtering non-canonical introns and allowing up to 10 mismatches per read and only keeping uniquely mapped reads. The genome index was constructed using the gene annotation supplied with the mm10 Illumina iGenomes collection and sjdbOverhang value of 100. Raw or FPKM (fragments per kilobase per million mapped reads) gene expression was quantified across all gene exons with HOMER v4.10.4 analyzeRepeats.pl with mm10 annotation v5.10 and parameters -strand + -count exons -condenseGenes (top-expressed isoform as proxy for gene expression), and differential gene expression was carried out on the raw counts with HOMER getDiffExpression.pl that runs DESeq2 v1.14.1 using replicates to compute within-group dispersion. BMP6 treatment samples were adjusted for mouse-specific batch effects using the - batch flag. Differentially expressed genes were defined as having FDR<0.05, FC>1.5, and FPKM>1, when comparing two experimental conditions.

Pathway analysis was performed using PANTHER v13, using the over-representation test with Fisher’s Exact test and false discovery rate corrected^86^.

### Immunocytochemistry

#### Staining astrocytes for cell-specific markers

After 7 days (for astrocytes that were not treated with additional protein factors) or 12 days (for astrocytes treated with either BMP6 or noggin in minimal growth medium), astrocytes were fixed with warm (37°C) 4% paraformaldehyde (EMS 50980487) for 20 minutes. Coverslips were washed three times with room temperature (RT) PBS and blocked and permeabilized for 30 minutes in 50% goat serum/50% antibody buffer (150 mM NaCl; 50 mM Tris; 100 mM L-lysine; 1% BSA; pH 7.4) and 0.5% Triton X-100 (Sigma T9284). Coverslips were washed once with PBS and incubated overnight at 4°C with primary antibodies against GFAP (1:1000, Millipore, MAB360), Aqp4 (1:1000, Sigma, A5971), GLAST (1:100, Miltenyi 130-095-822), NeuN (1:1000, Millipore, MAB377), Iba1 (1:1000, Wako, 016-20001), or NG2 (1:1000, Millipore, AB5320) in a solution of antibody buffer + 10% goat serum. The following day, cells were washed three times with PBS and incubated for 1-2 hours with appropriate secondary antibodies (goat anti-rabbit Alexafluor 594 (1:1000, Thermo Fisher Scientific A-11037), goat anti-mouse Alexafluor 488 (1:1000, Thermo Fisher Scientific A-11029), goat anti-mouse Alexafluor 594 (1:1000, Thermo Fisher Scientific A-11032)) in a solution of antibody buffer + 10% goat serum, before being washed three times and mounted with SlowFade + DAPI (Life Technologies s36939) on glass slides (Fisherfinest 12-244-2) and sealed with clear nail polish. Astrocytes were imaged using a Zeiss AXIO Imager Z2 motorized microscope (430000-9902) using a 20X 0.8NA objective. Images were analyzed using ImageJ by calculating the % of total cells (DAPI) that were positive for the cell marker.

#### Neurite staining of cortical neurons

After 48 hours in culture, cortical neurons were fixed for 10 minutes with 4% paraformaldehyde in PBS warmed to 37°C. Coverslips were washed three times with RT PBS and blocked and permeabilized for 30 minutes in 50% antibody buffer/50% goat serum and 0.5% Triton X-100. Coverslips were washed once with PBS and incubated overnight at 4°C with primary antibodies against MAP2 (to mark dendrites) (1:5000, EnCor Biotechnologies CPCA-MAP2) and Tau (to mark axons) (1:500, Millipore Sigma MAB3420) in a solution of antibody buffer + 10% goat serum. The following day, cells were washed three times with PBS and incubated for 1-2 hours with secondary antibodies (goat anti-chicken Alexafluor 488 (1:1000, Thermo Fisher Scientific A11039), goat anti-mouse Alexafluor 594 (1:1000)) in a solution of antibody buffer + 10% goat serum, before being washed three times with PBS and mounted with SlowFade + DAPI on glass slides, sealed with nail polish, and stored at −20°C until imaging. Cortical neurons were imaged using a 10X 0.45NA objective on a Zeiss AXIO Imager Z2 motorized microscope (430000-9902). For each assay, there were 3 coverslips per condition, and 20 images were taken per coverslip, for a total of 60 images per condition per assay with at least 1 neuron per image. Images were processed in ImageJ to merge all channels, convert the images to 16-bit, and apply a normalization of 0.1. Images were analyzed using MetaMorph software (Molecular Devices), using the Neurite Outgrowth Module to measure total, mean, max, and median neurite outgrowth length, number of processes per cell, and number of process branches per cell.

### Immunohistochemistry

#### Tissue collection and preparation

P7 male littermate mice were used for all experiments (Aldh1L1-eGFP;Mecp2-/y and Aldh1L1-eGFP;Mecp2+/y, or Aldh1L1-eGFP;Fmr1-/y and Aldh1L1-eGFP;Fmr1+/y). Mice were anesthetized with intraperitoneal injection of 100 mg/kg Ketamine (Victor Medical Company) and 20 mg/kg Xylazine (Anased) mix and transcardially perfused with PBS followed by 4% PFA. Brains were dissected and postfixed in 4% PFA at 4°C overnight. Brains were washed three times in PBS and cryoprotected in 30% sucrose at 4°C. After at least 48 hours in sucrose, brains were frozen in TFM (General data healthcare TFM-5) in a dry ice/ethanol mix and stored at −80°C until use. Fixed tissue was used for immunohistochemistry.

#### Immunohistochemical staining for Igfbp2 and pSMAD in P7 mouse brains

Coronal sections were collected on Superfrost Plus micro slides (VWR 48311-703) at 14 um thickness (3.4 mm posterior to Bregma) made on a cryostat (Hacker Industries OTF5000). Sections were blocked and permeabilized for 1 hr at room temperature in a humidified chamber in antibody buffer + 0.3% Triton X-100 in PBS. Primary antibodies, goat anti-Igfbp2 (1:500, R&D AF797) or rabbit anti-pSMAD (1:800, Cell Signaling 9516), were incubated in antibody buffer + 0.3% Triton X-100 in a humidified chamber overnight at 4°C. Slices were washed three times with PBS and incubated for 1-2 hr at room temperature with appropriate secondary antibody, donkey anti-goat Alexa 594 (1:500, Thermo Fisher Scientific A-11058) or goat anti-rabbit Alexa 594 (1:500, Thermo Fisher Scientific A-11037) in antibody buffer + 0.3% Triton X-100. Sections were washed with PBS three times and mounted with SlowFade gold antifade mountant with DAPI. Coverslips (22 mm x 50 mm 1.5 thickness (Fisher Scientific 12-544-D)) were placed on top of the sections and sealed with nail polish (Electron Microscopy Sciences 72180). Negative controls included sections where Igfbp2 or pSMAD primary antibodies were omitted.

For Igfbp2 layer 2/3 of the visual cortex was imaged using a Zeiss LSM 880 Rear Port Laser Scanning Confocal Microscope. The investigator was blinded to the genotypes of the mice during sectioning, staining, imaging, and analysis. Images were taken using a 63X objective, as 16 bit images, and as z stacks of 10 steps with a total thickness of 3.85um. Acquisition settings were the same for WT and KO samples imaged in the same experiment (littermate pairs). Imaging was performed on 3 sections (technical replicates) from 6 mice (biological replicates) per genotype. Analysis was performed using IMARIS software (Bitplane). Astrocytes (eGFP) were defined as surfaces with background subtraction (largest sphere contained within the surface set to 5um) and smoothed to 0.110um. Igfbp2 (channel 594) was masked within and without these surfaces to allow separate analysis of internal and external Igfbp2, which was also defined as surfaces. Representative images are snapshots of the 3D z-stack image (left), z-stack with internal Igfbp2 masked i.e. removed (middle), and 3D surfaces of astrocytes and extracellular Igfbp2 (right).

For pSMAD all layers of the visual cortex were imaged using a Zeiss LSM 880 Rear Port Laser Scanning Confocal Microscope. The investigator was blinded to the genotypes of the mice during sectioning, staining, imaging, and analysis. Images were acquired using a 20X objective, as 16 bit images of 4-6 tiles with 10% overlap, z stacks of 3 steps with a total thickness of 2.27 um. Imaging was performed on 3 sections (technical replicates) from 3 mice (biological replicates) per genotype. Images were used to generate maximum intensity projections and exported as TIFF images for analysis. Using ImageJ software (NIH), ROIs of 1800 x 1200 pixels were drawn in each image, capturing layers I-IV of the cortex. Expression of pSMAD in the nucleus was measured using the particle analysis tool in ImageJ and counted using the Cell Counter plugin (Kurt de Vos). DAPI was used to create a mask, and proportion of pSMAD positive astrocytes was determined by counting the number of colocalized DAPI + pSMAD + eGFP+ cells and normalizing to the total number of astrocytes.

### Statistical Analysis and Data Presentation

Statistical tests were performed using Prism 7. All tests are two-tailed. To compare more than two groups, one-way ANOVA with Dunnett’s post hoc test for multiple comparisons was used. Student’s t test was used to compare two groups. When data did not pass normal distribution test, multiple comparisons were done by Kruskal-Wallis ANOVA on ranks and pairwise comparisons were done with Mann-Whitney Rank Sum test. Exact p-values are reported on graphs or in the figure captions. Graphs and heatmaps were generated using Prism 7. Unless stated in the figure legend, all graphs represent average ± s.e.m.. For analysis of neurons in vitro, data is averaged from all cells and the number of cells indicated in the bar graph; for analysis of in vivo changes, data is averaged from each mouse, and data points on dot plots represent individual mice. Venn diagrams were generated using InteractiVenn http://www.interactivenn.net/

### Data Deposition

RNA sequencing raw data has been deposited in GEO, GSE139285. https://www.ncbi.nlm.nih.gov/geo/

The following secure token has been created to allow review of record GSE139285 while it remains in private status: sfezegusdtknbgx

The mass spectrometry proteomics data have been deposited to the ProteomeXchange Consortium via the PRIDE partner repository with the dataset identifier PXD015996. https://www.ebi.ac.uk/pride/archive/login

While the record is in private status it may be accessed by reviewers: Username; Password: NfVAguZZ

## Acknowledgements

We thank Cari Dowling for technical assistance, and A. Saghatelian, members of the Allen lab and MNL for helpful discussion. This work was supported by a Salk Institute Innovation grant. A.L.M.C. was supported by a Dennis Weatherstone Predoctoral Fellowship from Autism Speaks, and the Chapman Foundation. N.J.A. is supported by the Chan Zuckerberg Initiative, Hearst Foundation and Pew Foundation. Supported by core facilities of the Salk Institute including the Mass Spectrometry Core, Next Generation Sequencing Core, Integrative Genomics and Bioinformatics Core, Waitt Advanced Biophotonics Core, with funding from NIH-NCI CCSG: P30 014195, Chapman Foundation, Waitt Foundation and Helmsley Center for Genomic Medicine

## Author Contributions

A.L.M.C. and N.J.A. designed the experiments and wrote the manuscript with input from other authors. N.J.A. conceived the project. A.L.M.C. performed all experiments except mass spectrometry, performed by J.K.D.. N.J.A., A.L.M.C., J.K.D., and M.N.S. analyzed data.

## Supplemental Tables 1-7

All supplemental tables are Excel documents each consisting of data on multiple tabs.

**Supplemental Table 1 (related to Figure 2,6).** Proteomic profiles of astrocyte conditioned media from WT, RTT, FXS, DS, and BMP6-treated astrocytes in vitro. Spectral count data normalized as NSAF; a cut off was set at 0.01% of total protein to be included.

**Supplemental Table 2 (related to Figure 3,6,7).** Change in protein secretion between FXS, RTT, DS and BMP6-treated ACM compared to WT, as well as overlapping changes between all 3 NDs, and all 3 NDs and BMP6-treated ACM.

**Supplemental Table 3 (related to Figure 3,6).** Pathway analysis of protein secretion changes between FXS, RTT, DS and BMP6-treated ACM compared to WT.

**Supplemental Table 4 (related to Figure 2,6).** Gene expression profiles of WT, RTT, FXS, DS, and BMP6-treated astrocytes in vitro, presented as FPKM.

**Supplemental Table 5 (related to Figure 3,6,7).** Changes in gene expression between FXS, RTT, DS, and BMP6-treated astrocytes compared to WT, as well as all overlapping changes between all 3 NDs and BMP6-treated astrocytes compared to WT.

**Supplemental Table 6 (related to Figure 3,6).** Pathway analysis of gene expression changes in FXS, RTT, DS and BMP6-treated astrocytes compared to WT.

**Supplemental Table 7 (related to Figure 3,6).** Overlapping protein secretion and gene expression changes between FXS, RTT, DS and BMP6-treated astrocytes compared to WT.

