## Supplemental Figures for "Aberrant astrocyte protein secretion contributes to altered neuronal development in diverse disorders"

a

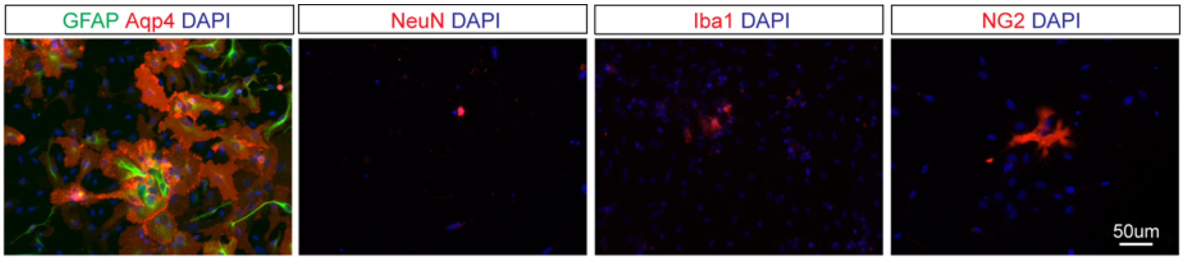


b % of DAPI+ cells expressing marker

| | Aqp4 | GFAP | NG2 | NeuN | Iba1 |
|---|---|---|---|---|---|
| Mean | 43.8 | 37.7 | 5.3 | 0.8 | 0.6 |
| SEM | 2.2 | 7.9 | 1.2 | 0.3 | 0.3 |

c

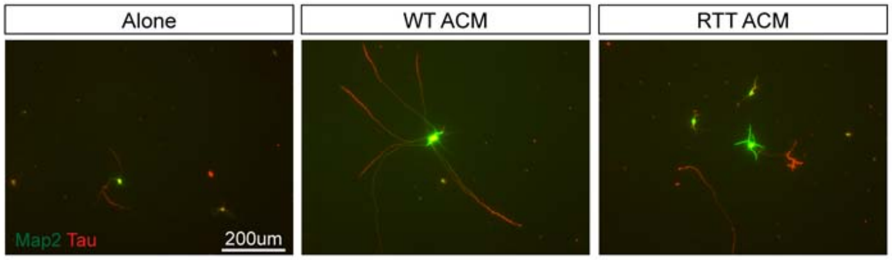


d

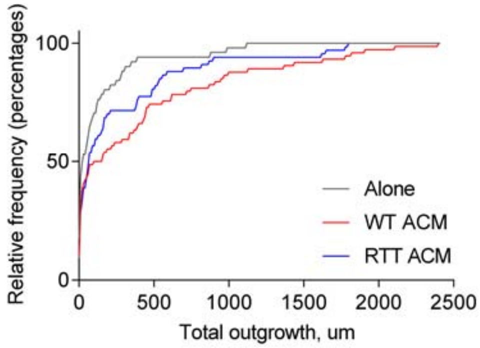


e

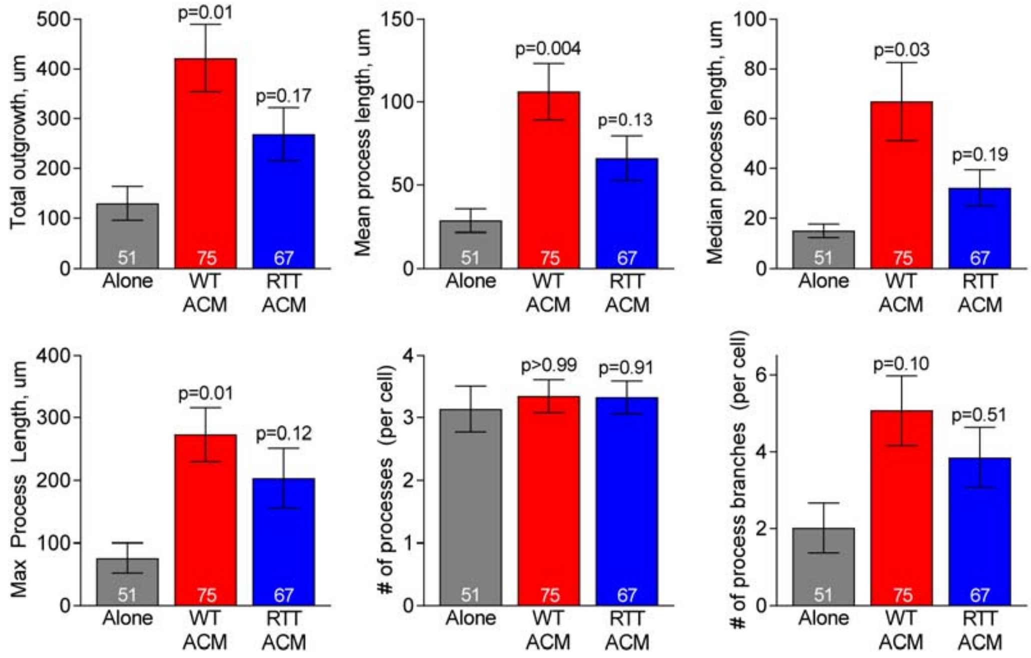


**Figure S1** (related to Figure 1). Immunopanned astrocyte and neuron cultures for the study of NDs.

**Figure S1 (related to Figure 1). Immunopanned astrocyte and neuron cultures for the study of NDs. a,b.** Immunostaining IP-astrocyte cultures for cell type markers reveals that the majority of cells express astrocyte-associated proteins Gfap and Aqp4, while rarely expressing NeuN (neuronal marker), Iba1 (microglial marker) or NG2 (oligodendrocyte precursor cell marker), N=15 astrocyte cultures (3WT, 1 RTT, 3 FXS, 8 DS; no differences were observed between ND and WT expression of cell markers). **c.** Example images from Figure 1e, prior to processing and analysis. WT neurons immunostained with MAP2 (dendrites, green) and tau (axon, red). **d.** Relative frequency of total neurite outgrowth, example experiment shown, same data as Figure 1f. **e.** Examination of additional measures of neurite growth for experiments in Figure 1e,f. Bar graphs mean±s.e.m. Number inside bar = number of neurons. Statistics by one-way ANOVA on ranks, p-values compared to neurons alone.

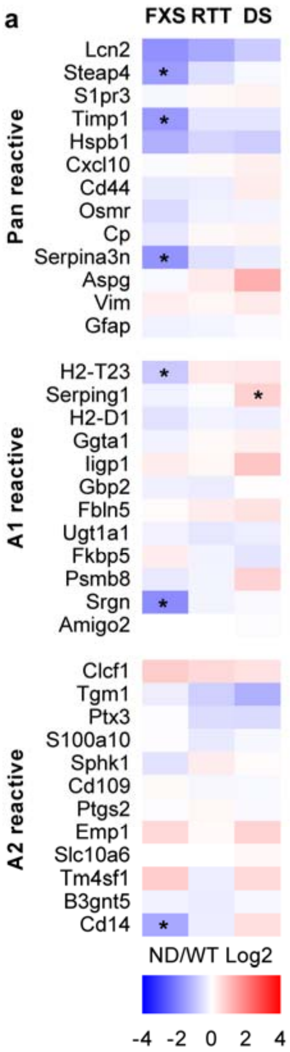


**Figure S2** (related to Figure 2). Immunopanned astrocytes reproduce known in vivo alterations to ND astrocyte function.

**Figure S2 (related to Figure 2). Immunopanned astrocytes reproduce known alterations to ND astrocyte function. a.** Reactive astrocyte markers from pan reactive, A1 reactive (inflammatory) and A2 reactive (supportive) astrocytes are not consistently altered in IP astrocyte cultures from ND compared to WT, demonstrating cultures are not reactive. Data from RNA sequencing. N=6 cultures WT, RTT, FXS; 4 DS. * adjusted $p<0.05$, FPKM>1 and fold change ≥1.5.

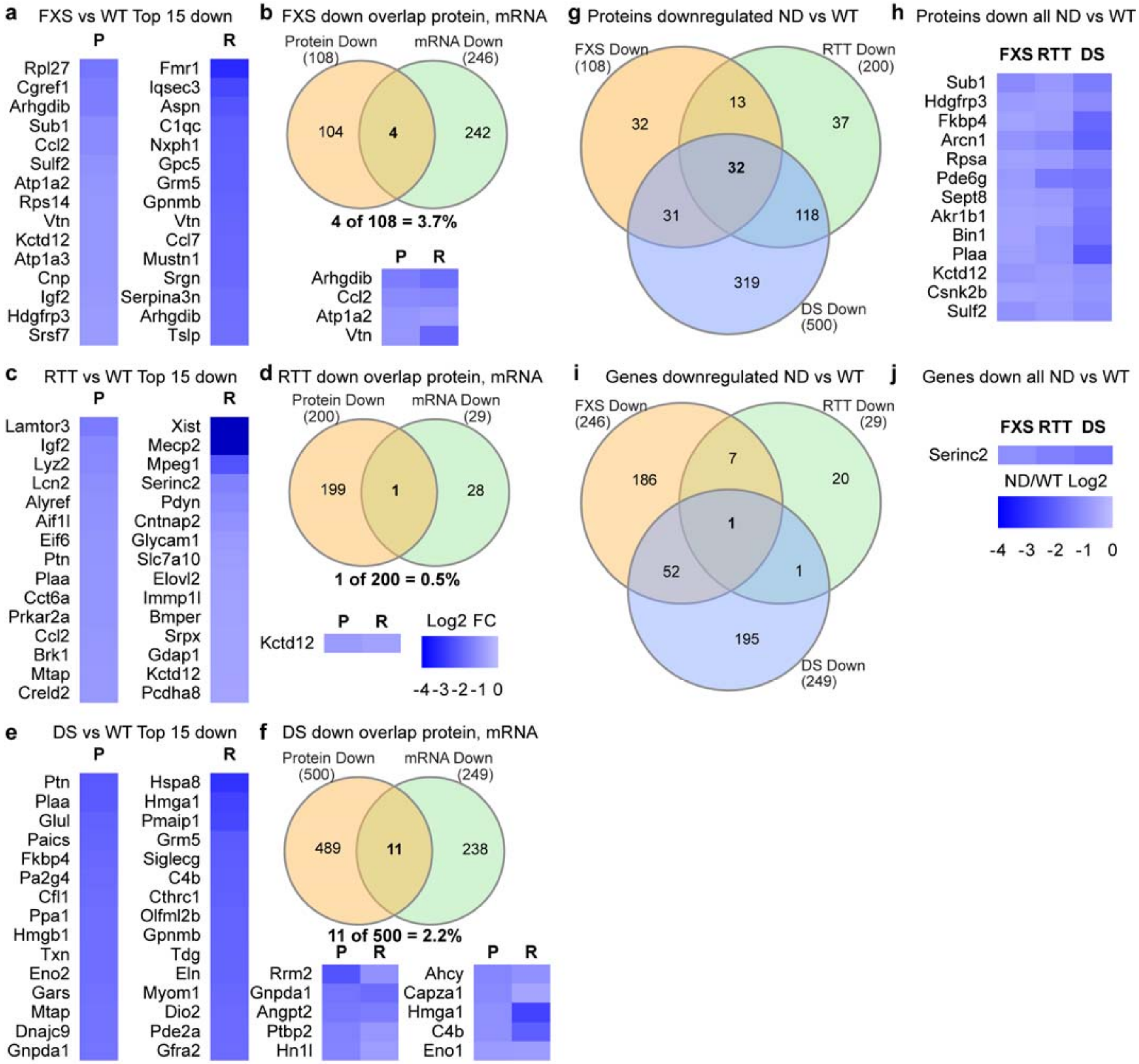


**Figure S3** (related to Figure 3). Identification of altered protein secretion and gene expression between astrocytes from WT and NDs.

**Figure S3 (related to Figure 3). Identification of altered protein secretion and gene expression between astrocytes from WT and NDs. a,c,e.** Heatmaps of top-fold change in proteins (P) and genes (R) decreased between WT and FXS (**a**), RTT (**c**) and DS (**e**) ACM and astrocytes. **b,d,f.** Venn diagram of overlap between proteins and genes with decreased level in FXS (**b**), RTT (**d**) and DS (**f**) ACM and astrocytes **g,h.** Venn diagram showing overlap in proteins decreased in all ND (**g**), and heatmap of altered proteins ranked by abundance in WT ACM (**h**). **i,j.** Venn diagram showing number of genes decreased (**i**) and corresponding heatmap (**j**) of overlapping altered genes. Scale bar in **d** applies to all Heatmaps. Proteomics, N=6 cultures per genotype, *$p<0.05$, abundance >0.01%, fold change between WT and ND ≥1.5. RNASeq, N=6 cultures WT, RTT, FXS; 4 DS, * adjusted $p<0.05$, FPKM>1, fold change between ND and WT ≥1.5.

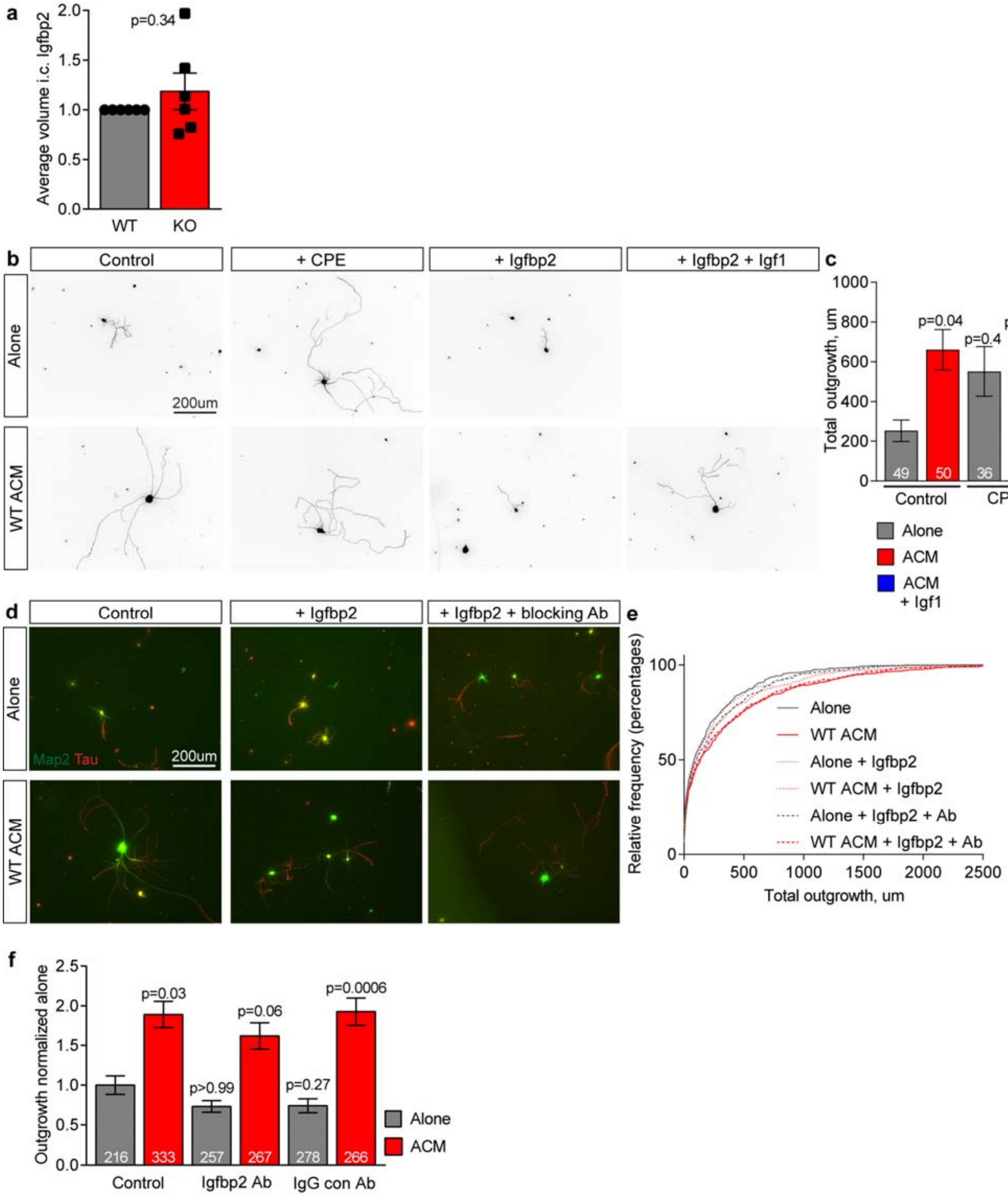


**Figure S4** (related to Figure 4). Excess Igfbp2 in ACM inhibits neurite outgrowth.

**Figure S4 (related to Figure 4). Excess Igfbp2 in ACM inhibits neurite outgrowth. a.** Immunostaining for Igfbp2 in RTT and WT P7 cortex shows no changes in intracellular Igfbp2. N=6 WT, 6 RTT mice. Bar graph mean±s.e.m. individual data points mice; statistics by T-test. **b,c.** Addition of Igfbp2 protein to WT ACM inhibits WT neurite outgrowth, which is reversed by adding IGF1. Addition of CPE protein to WT ACM does not inhibit WT neurite outgrowth. **b.** Example images of WT neurons cultured for 48 hours, conditions as marked. **c.** Quantification of total neurite outgrowth. Example experiment shown, repeated 2 times with same result. **d.** Example images from Figure 4g prior to processing and analysis. Neurons immunostained with MAP2 (dendrites, green) and tau (axon, red). **e**. Relative frequency distribution plot of total neurite outgrowth length, pooled data from 3 experiments, same as Figure 4h. **f.** Quantification of total neurite outgrowth determined that the Igfbp2 Ab or an IgG Control antibody had no effect on neurite outgrowth alone or with WT ACM. Example experiment shown, repeated twice with same result. Bar graphs (c,f) represent mean±s.e.m. Number inside bar = number of neurons. Statistics by one-way ANOVA on ranks, p values compared to control alone condition.

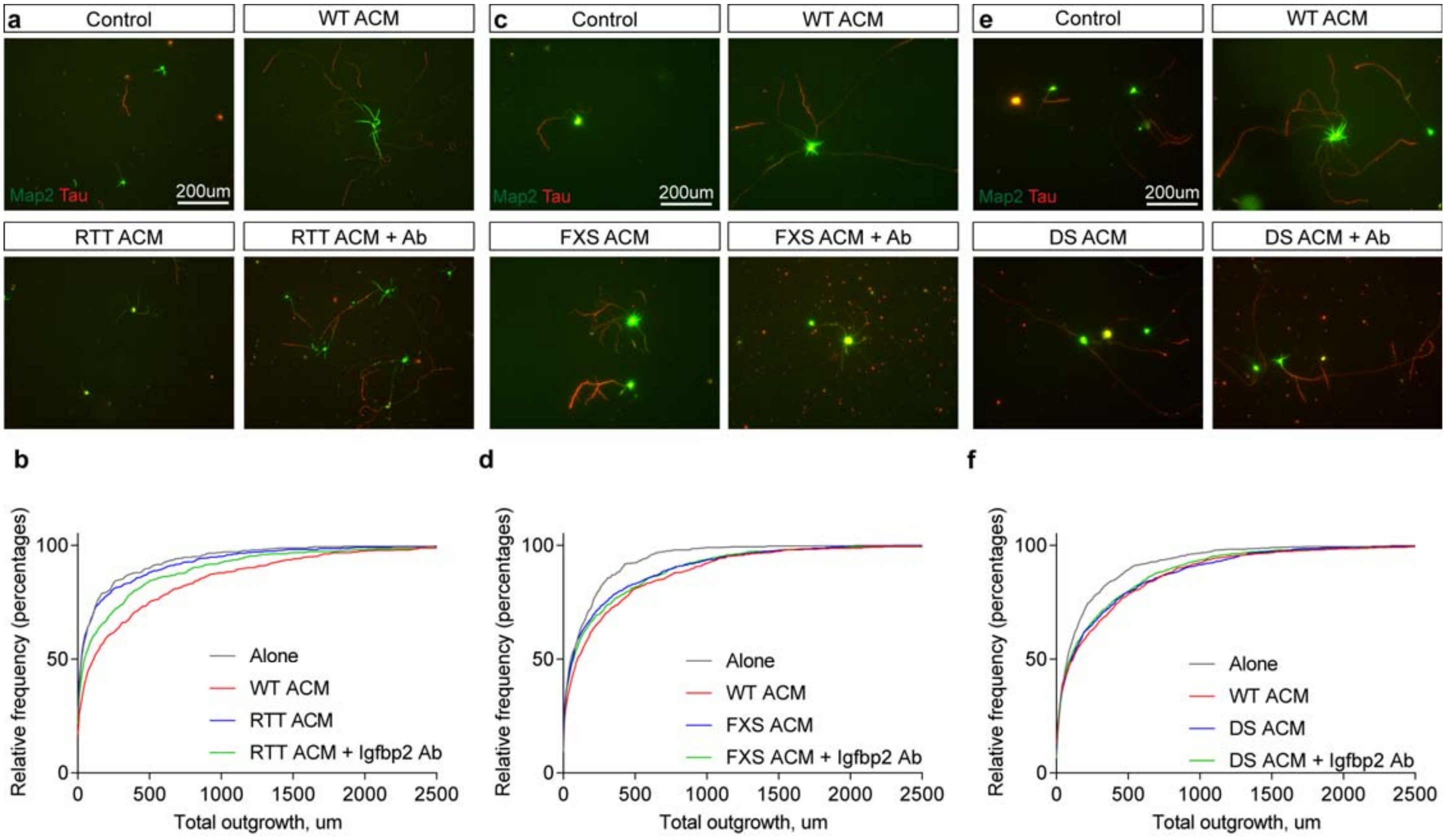


**Figure S5** (related to Figure 5). Blocking Igfbp2 in RTT ACM reduces neurite outgrowth deficits.

**Figure S5 (related to Figure 5). Blocking Igfbp2 in RTT ACM reduces neurite outgrowth deficits. a,c,e.** Example images from Figure 5a,c,e prior to processing and analysis. Neurons cultured for 48 hours in RTT (**a**), FXS (**c**) or DS (**e**) ACM and immunostained with MAP2 (dendrites, green) and tau (axon, red). **b,d,f**. Relative frequency distribution plot of total neurite outgrowth, pooled data from 3 (b), 4 (d), 5 (f) experiments per graph, same data as Figure 5b,d,f.

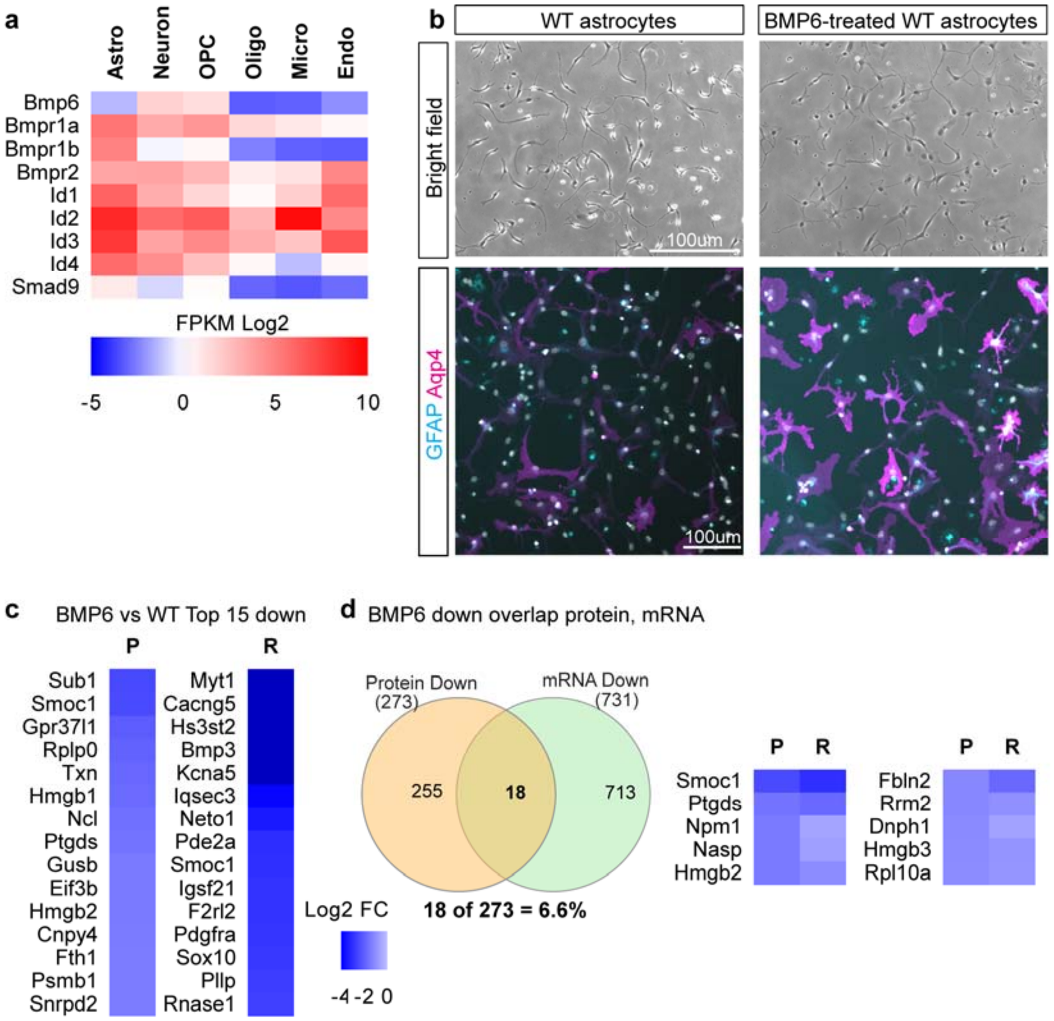


**Figure S6** (related to Figure 6). Activating BMP signaling in astrocytes induces changes that overlap with ND astrocytes.

**Figure S6 (related to Figure 6). Activating BMP signaling in astrocytes induces changes that overlap with ND astrocytes. a.** Relative expression of BMP family members in purified cell types in the cortex shows enrichment for BMP target genes in astrocytes (data from Zhang et al, 2014). **b**. BMP6-treated astrocytes have thinner more branched processes (top), and increased expression of both GFAP (cyan) and AQP4 (magenta) (bottom). Example images shown, experiment repeated 3 times with same effect. **c.** Heatmaps of proteins (P) and mRNA (R) decreased between BMP6-treated and untreated WT ACM and astrocytes, ranked by fold-change. **d.** Venn diagram showing overlap between proteins and genes with decreased expression in WT astrocytes treated with BMP6. For mass spectrometry and RNA Sequencing N=6 cultures, half of each culture treated with BMP6 and other half left untreated. Proteomics, *$p<0.05$, abundance >0.01%, fold change ≥1.5. RNASeq, * adjusted $p<0.05$, FPKM>1, fold change ≥1.5.

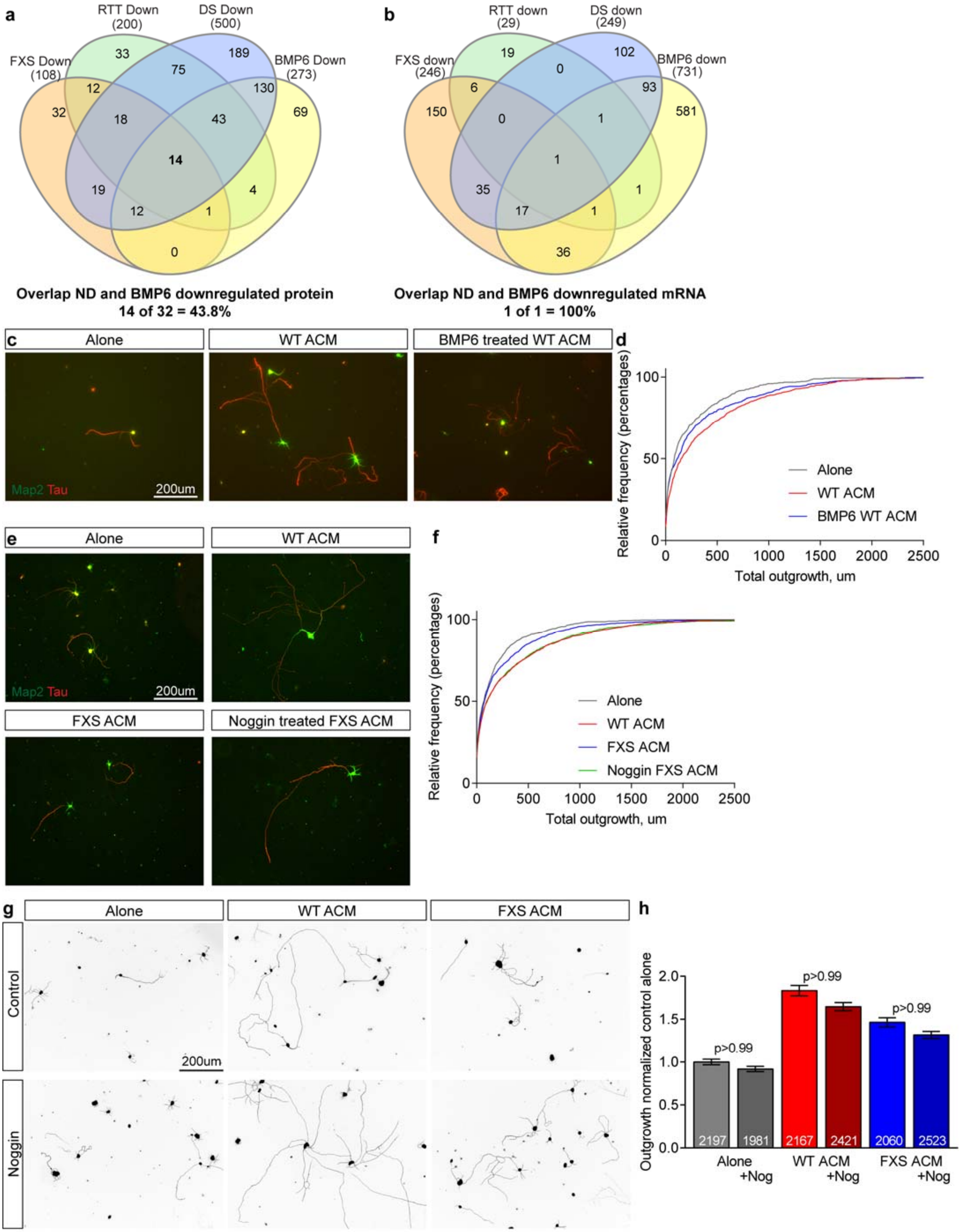


**Figure S7** (related to Figure 7). Blocking Bmp signaling in FXS astrocytes abolishes neurite outgrowth deficits.

**Figure S7 (related to Figure 7). Blocking BMP signaling in FXS astrocytes abolishes neurite outgrowth deficits. a,b.** BMP6-treated WT astrocytes show protein secretion (**a**) and gene expression (**b**) decreases that overlap with ND astrocytes. N=6 cultures each WT, FXS, RTT, DS, plus 6 cultures WT +/- BMP6. **c.** Example images from Figure 7d prior to processing and analysis. Neurons immunostained with MAP2 (dendrites, green) and tau (axon, red). **d**. Relative frequency distribution plot of total neurite outgrowth length, same data as Figure 7e. **e.** Example images from Figure 7g prior to processing and analysis. Neurons immunostained with MAP2 (dendrites, green) and tau (axon, red). **f**. Relative frequency distribution plot of total neurite outgrowth, same data as Figure 7h. **g**. Example images of cortical neurons treated with noggin at the time of plating, ± WT ACM or ± FXS ACM. **h**. Quantification of total neurite outgrowth. Bar graph mean±s.e.m. Number inside bar = number of neurons, pooled from 3 experiments for all graphs. Statistics by one-way ANOVA on ranks, p-values against paired condition i.e. plus/minus noggin.
